# Single-Cell RNA-Seq Reveals Endocardial Defect in Hypoplastic Left Heart Syndrome

**DOI:** 10.1101/809731

**Authors:** Yifei Miao, Lei Tian, Marcy Martin, Sharon L. Paige, Francisco X. Galdos, Jibiao Li, Alyssa Guttman, Yuning Wei, Jan-Renier Moonen, Hao Zhang, Ning Ma, Bing Zhang, Paul Grossfeld, Seema Mital, David Chitayat, Joseph C. Wu, Marlene Rabinovitch, Timothy J. Nelson, Shuyi Nie, Sean M. Wu, Mingxia Gu

**Affiliations:** Department of Pediatrics, Division of Pediatric Cardiology; Vera Moulton Wall Center for Pulmonary Vascular Disease; Stanford Cardiovascular Institute; Institute of Stem Cell and Regenerative Biology, Stanford School of Medicine, Stanford, California, USA; School of Biological Sciences, Georgia Institute of Technology, Atlanta, Georgia, 30332, USA; Center for Personal Dynamic Regulomes, Stanford School of Medicine, Stanford, California, 94305, USA; Key Laboratory of Systems Biomedicine, Shanghai Center for Systems Biomedicine, Xin Hua Hospital, Shanghai Jiao Tong University, Shanghai 200240, China; Department of Pediatrics, UCSD School of Medicine, La Jolla, California, 92093, USA; Department of Pediatrics, Hospital for Sick Children, University of Toronto, Toronto, Ontario M5G 1X8, Canada; The Prenatal Diagnosis and Medical Genetics Program, Department of Obstetrics and Gynecology, Mount Sinai Hospital, University of Toronto, Toronto, Ontario M5G 1X5, Canada; Division of General Internal Medicine, and Pediatric Cardiology, and Department of Molecular Pharmacology and Experimental Therapeutics, Mayo Clinic, Rochester, Minnesota, 55905, USA; Co first author; Lead Contact

**Keywords:** Hypoplastic left heart syndrome, Single-cell RNA-seq, Induced pluripotent stem cells, Human heart tissue, Endocardium, Endothelial to mesenchymal transition, NOTCH, Fibronectin, *De novo* mutation, ETS1, CHD7

## Abstract

Hypoplastic left heart syndrome (HLHS) is one of the most challenging forms of congenital heart diseases. Previous studies were mainly focused on intrinsic defects in myocardium. However, this does not sufficiently explain the abnormal development of the cardiac valve, septum, and vasculature, known to originate from the endocardium. Here, using single-cell RNA profiling, induced pluripotent stem cells, and human fetal heart tissue with an underdeveloped left ventricle, we identified a developmentally impaired endocardial population in HLHS. The intrinsic endocardial deficits contributed to abnormal endothelial to mesenchymal transition, NOTCH signaling, and extracellular matrix organization, all of which are key factors in valve formation. Consequently, endocardial abnormalities conferred reduced proliferation and maturation of cardiomyocytes through a disrupted fibronectin-integrin interaction. Several recently described HLHS *de novo* mutations were associated with abnormal endocardial gene and *FN1* regulation and expression. Our studies provide a rationale for considering endocardial function in future regenerative strategies for HLHS.

## Introduction

Hypoplastic left heart syndrome (HLHS) is a congenital heart malformation characterized by severe underdevelopment of the left ventricle, mitral valve, aortic valve, and ascending aorta. Previous studies using human induced pluripotent stem cells (iPSCs) from HLHS patients as well as mouse models of HLHS have shown an intrinsic defect in cardiomyocytes, with reduced cardiac differentiation efficiency, disorganized sarcomeres, abnormal mitochondria, and impaired NOTCH signaling (Liu et al., 2017; Yang et al., 2017). However, none of these features sufficiently explained the abnormal development of the cardiac valve, septum, and vasculature in HLHS.

Endocardial cells are specialized endothelial cells (EC) that form the innermost layer of the heart wall. The endocardium plays several pivotal roles in heart development and disease(Zhang et al., 2018). It serves as the source of mesenchymal cells in the endocardial cushions that give rise to the structural elements of the atrioventicular valves as well as the atrial and membranous ventricular septa(Harris and Black, 2010). The “no flow, no grow” theory suggests that the abnormal cardiac valve could limit blood flow, leading to the ventricular hypoplasia(Boselli et al., 2015). Endocardial cells also contribute to a small amount of coronary vascular ECs in the developing heart(Tian et al., 2015). Moreover, the reciprocal endocardial-myocardial interaction constitutes an important signaling center that mediates the trabeculation and maturation of the heart chamber(Bressan et al., 2014; de la Pompa and Epstein, 2012). Given that the majority of congenital heart defects are represented by valvular abnormalities, septal defects, and/or ventricular underdevelopment(Botto et al., 2001; Hoffman and Kaplan, 2002), it is important to understand the role of the endocardium in the pathogenesis of these heart diseases. This will also inform the development of therapeutic strategies aimed at ventricular and valvular regeneration. Previously, it has been reported that HLHS patient hearts had a reduced EC population and lower capillary density compared with normal hearts(Gaber et al., 2013). HLHS fetal heart ECs also exhibited the greatest susceptibility to genotoxic injury compared with other cardiac cell types(Gaber et al., 2013). Several genetic variants (e.g., in *NOTCH1*, *HAND1*) that are robust determinants of endocardial and endothelial development and function(Luxan et al., 2016; Theodoris et al., 2015), are also implicated in HLHS(Hrstka et al., 2017; Kobayashi et al., 2014; Yang et al., 2017). However, the nature of endocardial insufficiency as an underlying mechanism causing HLHS remains poorly understood.

To elucidate the transcriptomic and functional defects in the HLHS endocardium, we carried out single-cell RNA sequencing (scRNA-seq) in both HLHS iPSC-ECs and fetal heart tissue with an underdeveloped left ventricle. We identified a developmentally impaired endocardial cell population in HLHS compared with controls. Subsequent functional assays revealed that the endocardial defects in HLHS patient heart could lead to impaired endocardial to mesenchymal transition and cardiac valve formation. Additionally, our studies revealed that endocardial abnormalities in HLHS patients could contribute to the reduced proliferation and maturation of cardiomyocytes (CMs), mediated by a disrupted fibronectin-integrin α5β1 interaction. These features were dysregulated by reduced expression of the transcription factor ETS proto-oncogene 1 (ETS1) and other epigenetic regulators such as CHD7, which have been previously reported to have a *de novo* mutation associated with a hypoplastic left ventricle(Jin et al., 2017). Taken together, our findings have uncovered a mechanism whereby a developmentally impaired endocardium can underlie the ventricular and valvular hypoplasia in HLHS.

## Results

### HLHS patient *de novo* mutations were highly enriched in the endocardial and endothelial populations in a developing human heart

It is well appreciated that the genetics of HLHS is complex. Previously, a set of congenital heart disease (CHD) related *de novo* mutations (DNMs) were reported(Jin et al., 2017), some of which were strongly associated with HLHS. To further understand the role of these HLHS-associated DNMs in the development of human heart, we first examined their expression level at single cell resolution. A normal human fetal heart at a gestational age of 83 days was micro-dissected into the left- and right-side of the heart. Each side was enzymatically dispersed as single cells, enriched with endocardial/endothelial cells using a CD144 antibody and processed for single-cell RNA-sequencing (scRNA-seq) (Figure 1A). The majority of the cell types expected to be present in the heart were successfully retrieved by scRNA-seq analysis as shown in the UMAP projection, including a certain amount of CMs (*TNNI3^+^*) in addition to pan-ECs (*CDH5^+^*) (Figure 1B & C). Of note, *CDH5*^+^ populations clustered into three groups: endocardium (*NPR3^+^*), endothelium (*APLN^+^*) (unless specified, endothelium refers to the vascular endothelium), and lymphatic endothelium (*LYVE1^+^*) (Figure 1C).

**Figure 1.**
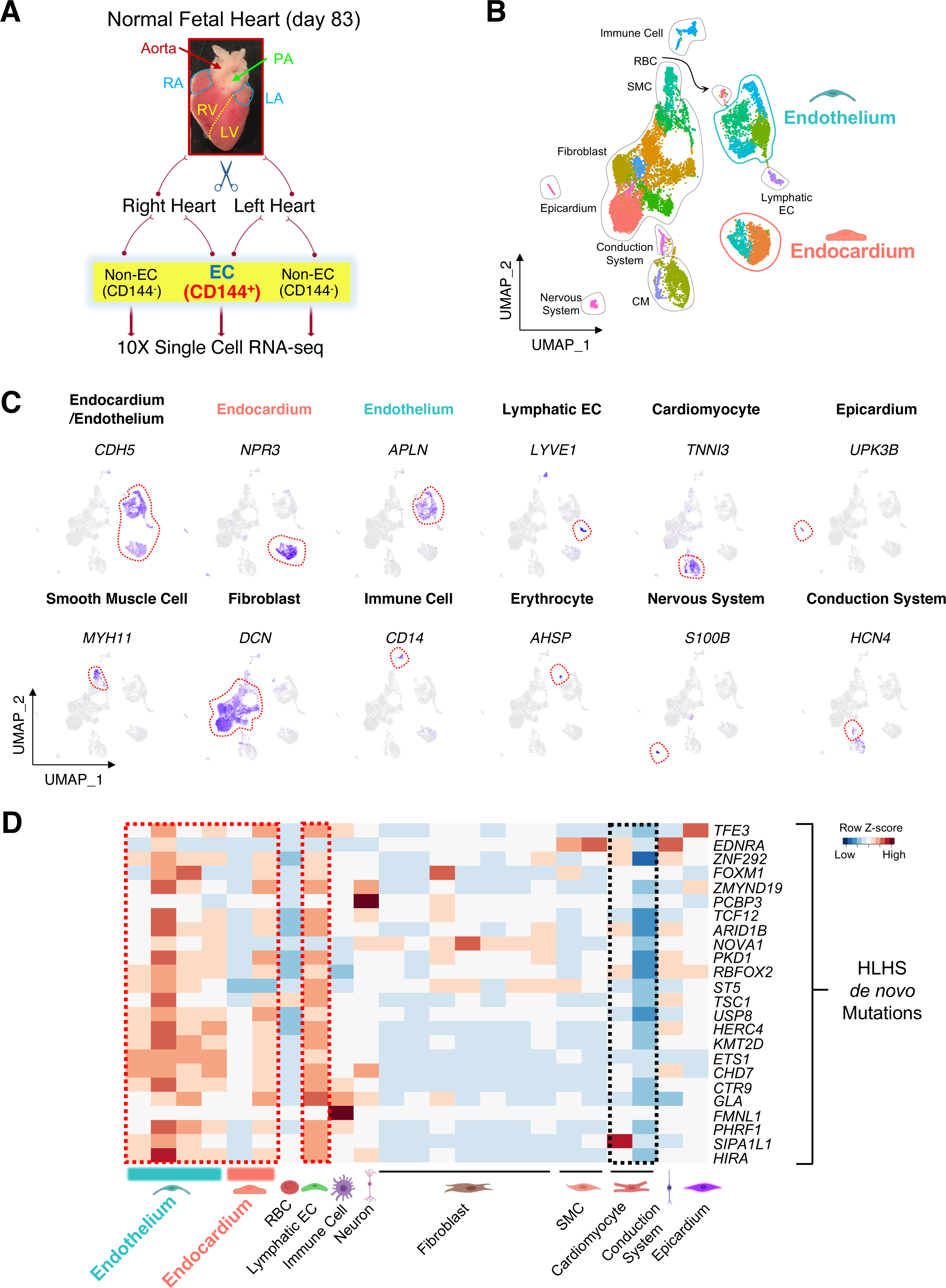
HLHS *de novo* mutations (DNMs) were enriched in the endocardium and endothelium based on scRNA-seq analysis of human fetal heart. (A) Schematic of the workflow for micro-dissection of normal human fetal heart and scRNA-seq. Endocardial/endothelial populations were enriched with a CD144 antibody and Magnetic-Activated Cell Sorting. PA: pulmonary artery; RA: right atrium; LA: left atrium; RV: right ventricle; LV: left ventricle. (B) UMAP projection of various cell types from day 83 normal human fetal heart. SMC: smooth muscle cell; RBC: red blood cell; CM: cardiomyocyte; EC: endothelial cell. (C) UMAP projection of represented genes for various cell types colored by represented genes’ expression level (purple indicates high expression level) in human fetal heart. (D) Cell type-specific RNA expression of HLHS DNM genes based on scRNA-seq from day 83 normal fetal heart. Row Z-score indicates RNA level. Red denotes high expression, blue minimal expression. See also Figure S1.

An in-depth analysis of CD144^+^ populations from the fetal heart further defined several endocardial and endothelial subpopulations (Figure S1A). The homologues of several well-defined genes that label mouse endocardium(Zhang et al., 2016), such as *NPR3*, *CDH11*, *HAPLN1*, and *ADGRG6* were also enriched in the human heart endocardium (Figure S1B). A small cluster of cells that were in close proximity to the endocardium highly expressed valve markers (*FST* and *PROX1*) (Figure S1C)(Hulin et al., 2019; Su et al., 2018). Within the cardiac vascular endothelial populations, there were distinct clusters expressing artery, vein, and capillary markers(Su et al., 2018; Wolf et al., 2019) as well as intermediate clusters that were positive for both arterial and venous/capillary genes (Figure S1D & E).

After generating the fetal heart atlas at single cell resolution, we next examined cell-type specific mRNA expression levels of the HLHS-associated DNMs, which have been reported previously(Jin et al., 2017). Intriguingly, the majority of these genes were highly expressed in the endocardium, endothelium, and lymphatic ECs (Figure 1D, red dashed box), as opposed to the cardiomyocytes (Figure 1D, black dashed box). These observations are consistent with a pivotal role for endocardial/endothelial dysfunctions in the pathogenesis of HLHS.

### scRNA-seq of HLHS iPSC-ECs uncovered endocardial defects

To further investigate the role of EC dysfunction in causing HLHS, iPSCs from four controls and three HLHS patients were differentiated into heterogenous ECs including an endocardial population using a previously established protocol(Gu, 2018) (Figure 2A). iPSC lines from HLHS cases that developed early-onset right heart failure, as defined in Table 1 were used in this study. After purification using magnetic sorting with an antibody to the EC surface marker CD144, day 8 iPSC-derived ECs (iPSC-ECs) expressed both endocardial and endothelial markers as compared to Day 0 undifferentiated iPSCs (Figure S2A). scRNA-seq of iPSC-ECs from control and HLHS patient revealed several EC subclusters (Figure 2B & C). Using the same endocardial markers identified in Figure 1 and Figure S2A, we found that control and HLHS patients were segregated with the control group represented by four clusters (2, 0, 5, and 3) and the HLHS cells by three clusters (6, 4, and 1). Control iPSC-ECs were highly enriched with endocardial genes (clusters 2 & 0); whereas HLHS iPSC-ECs showed a significantly reduced endocardial gene expression within only one small cluster (cluster 6) (Figure 2D-F). This finding was further validated by qPCR in 4 control and 3 HLHS iPSC-ECs (Figure 2G). Another finding was that HLHS iPSC-ECs demonstrated an arterial-to-venous phenotypic shift with a higher proportion of vein-like ECs rather than arterial ECs compared to the controls (Figure S2B-D).

**Figure 2.**
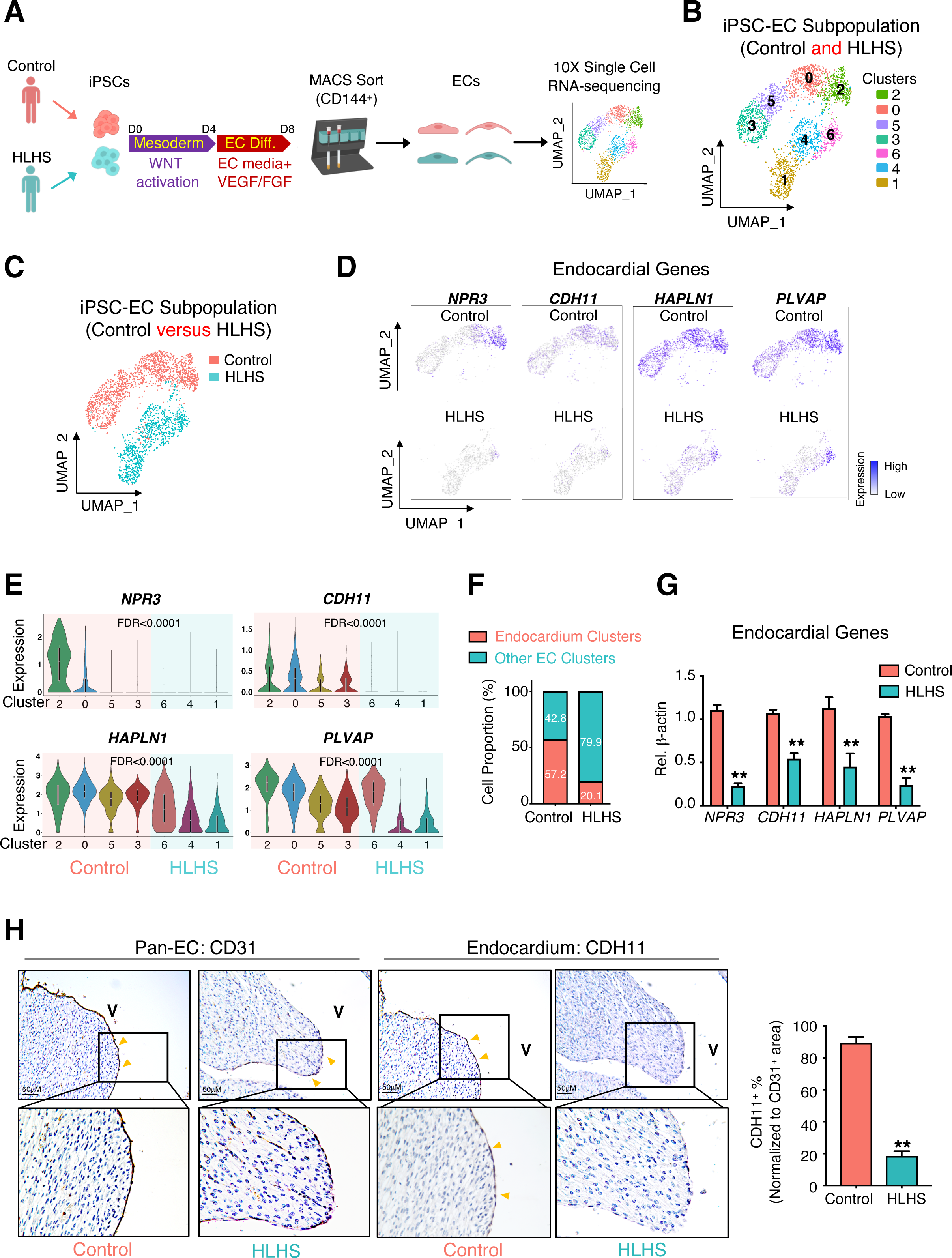
scRNA-seq of iPSC-ECs unraveled endocardial defects in HLHS patients. (A) Illustration of 10X genomics scRNA-seq of iPSC-ECs from one healthy control and one HLHS patient. (B) UMAP projection of iPSC-ECs from both the control and HLHS patient. Each color defines one subpopulation based on the transcriptomic phenotype. (C) UMAP projection colored by cell origin. (D) UMAP plots of iPSC-ECs colored by expression levels of endocardial markers. Purple denotes high expression, white minimal expression. (E) Violin plot visualization of endocardial gene expression distribution across different subpopulations. False discover rate (FDR) indicated the significant of difference between control (cluster 2, 0, 5, and 3) vs HLHS (cluster 6, 4, and 1). (F) Proportional distribution of endocardial clusters in iPSC-ECs based on scRNA-seq data. (G) Confirmation of endocardial genes expression changes between control (n=4) and HLHS (n=3) iPSC-ECs by quantitative PCR (qPCR). (H) Immunostaining of pan-EC (CD31) and endocardial (CDH11) proteins in fetal hearts from control (n=6) and HLHS (n=12) patients. In (G) and (H), data shown as the mean ± SEM. **p<0.01, control vs HLHS. See also Figure S2.

**Table 1.**
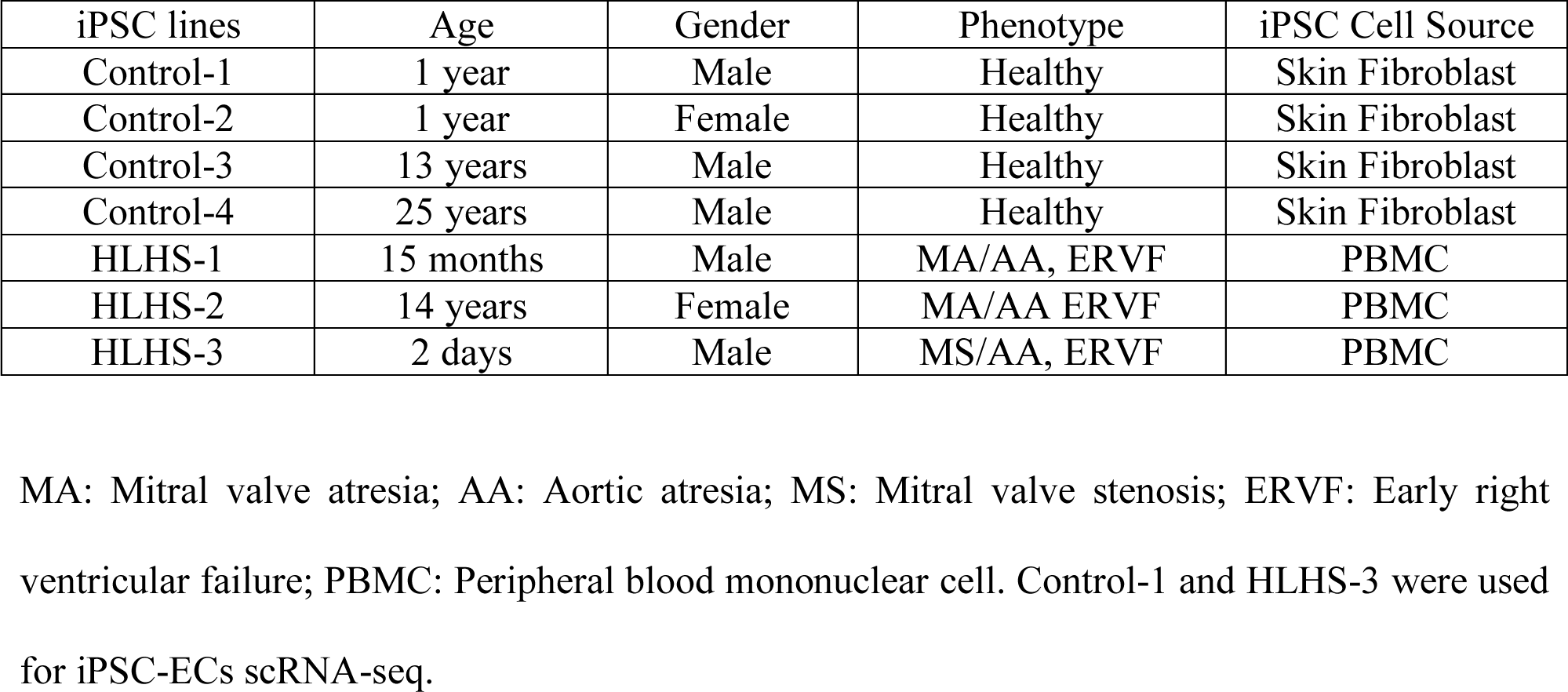
iPSC lines information

| iPSC lines | Age | Gender | Phenotype | iPSC Cell Source |
| --- | --- | --- | --- | --- |
| Control-1 | 1 year | Male | Healthy | Skin Fibroblast |
| Control-2 | 1 year | Female | Healthy | Skin Fibroblast |
| Control-3 | 13 years | Male | Healthy | Skin Fibroblast |
| Control-4 | 25 years | Male | Healthy | Skin Fibroblast |
| HLHS-1 | 15 months | Male | MA/AA, ERVF | PBMC |
| HLHS-2 | 14 years | Female | MA/AA ERVF | PBMC |
| HLHS-3 | 2 days | Male | MS/AA, ERVF | PBMC |

MA: Mitral valve atresia; AA: Aortic atresia; MS: Mitral valve stenosis; ERVF: Early right ventricular failure; PBMC: Peripheral blood mononuclear cell. Control-1 and HLHS-3 were used for iPSC-ECs scRNA-seq.

The abnormality in endocardial gene expression was also evident in HLHS patient heart tissues, where a reduction in the endocardial marker CDH11 expression was observed (Figure 2H, Figure S2E & F). A CD31 antibody clearly labeled the endocardium, coronary endothelium, and capillary endothelium (Figure S2E), while CDH11 was restricted to the endocardium, but not the coronary endothelium (Figure S2F). In control samples (n=6), the majority of the endocardial layer was positive for both CD31 and CDH11. However, CDH11 positive staining in the endocardium of HLHS heart ventricles (n=12) was significantly reduced (Figure 2H).

### HLHS iPSC-ECs exerted functional abnormalities in the endocardium

Based on the scRNA-seq results from iPSC-ECs, gene ontology (GO) enrichment analysis on endocardial clusters (clusters 2 and 0 in control; cluster 6 in HLHS) revealed suppression of several signaling pathways strongly relevant to endocardial functions as well as valvular structural remodeling and development, such as extracellular matrix (ECM) deposition, immune system, VEGF signaling and NOTCH signaling (Figure 3A) (Combs and Yutzey, 2009; Hulin et al., 2019; MacGrogan et al., 2010; Schroeder et al., 2003). ECM deposition is closely related to the endocardial to mesenchymal transition (EndoMT) (Brickner et al., 2000; Eisenberg and Markwald, 1995), which is a critical process for the development of valvular structures. Thus, we stimulated EndoMT in control and HLHS iPSC-ECs using TGF*β*2, and found that the protein level of the mesenchymal cell marker *α*-smooth muscle actin (*α*SMA) (Figure 3B) as well as EndoMT-related genes (Figure 3C) were significantly reduced in HLHS iPSC-ECs compared with control cells.

**Figure 3.**
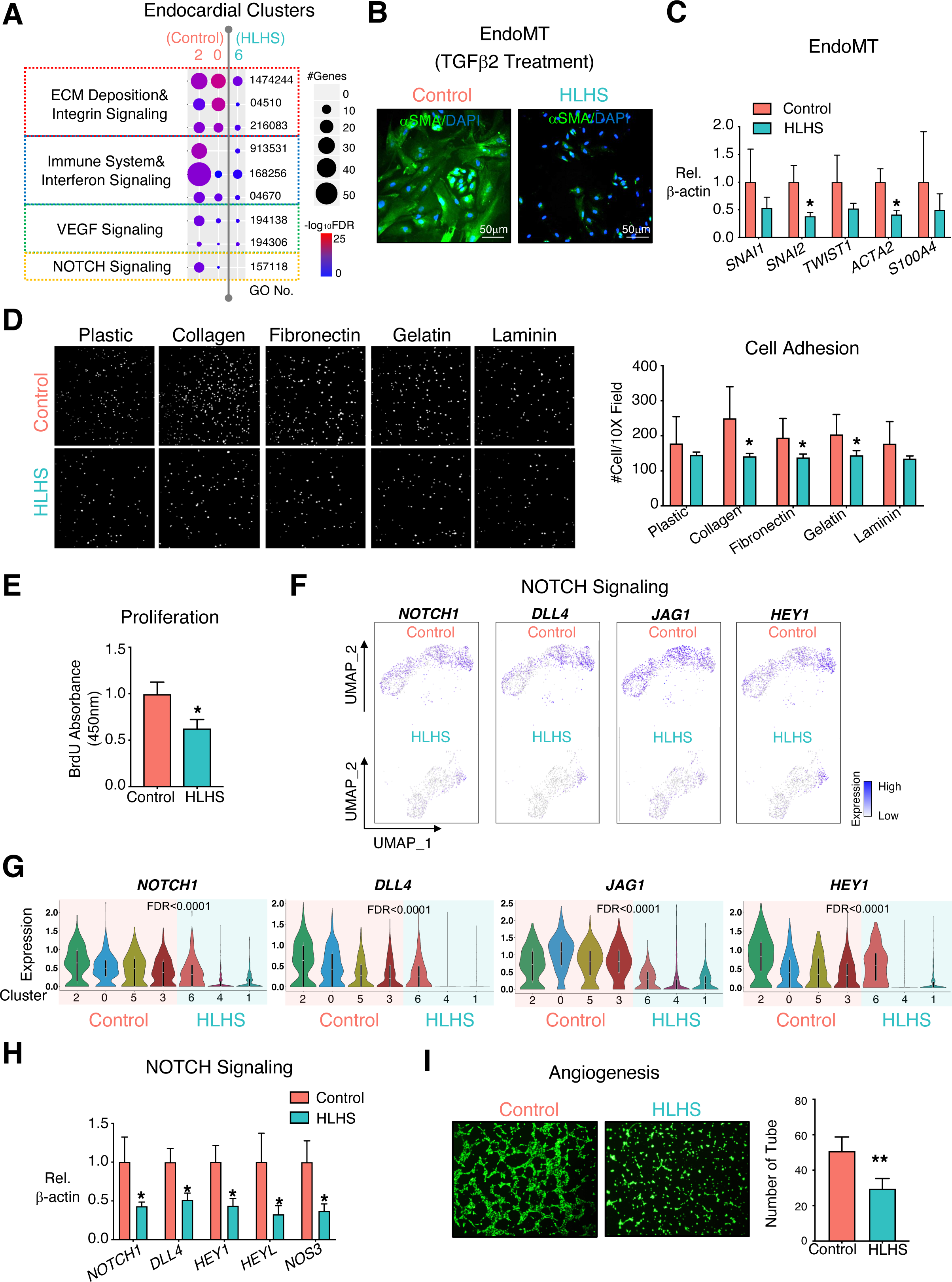
iPSC-ECs revealed functional defects in the HLHS endocardium. (A) Functional enrichment in endocardial clusters from control (cluster 2 & 0) and HLHS (cluster 6) iPSC-EC scRNA-seq. −log10FDR indicates the significance of enrichment. GO: gene ontology. *α* smooth muscle actin (*α*SMA) staining (B) and EndoMT related gene expression (C) in iPSC-ECs from control and HLHS patients after 7 days treatment with TGF*β*2 (10ng/μl). DAPI: nucleus. (D) Adhesion assay of iPSC-ECs with different ECMs to coat the culture dish. Each dot (DAPI nuclear staining) defines one single cell that was counted from four different views under each condition. Pictures are under 10X magnification. (E) Cell proliferation measurement in iPSC-ECs after 24-hour BrdU incorporation. UMAP (F) and violin plots (G) of the represented genes of the NOTCH pathway from scRNA-seq. FDR indicated the significance of difference between control (cluster 2, 0, 5, and 3) vs HLHS (cluster 6, 4, and 1). (H) Expression levels of NOTCH pathway related genes were measured by qPCR in control and HLHS patient iPSC-ECs. (I) Tube formation of control and HLHS iPSC-ECs after seeding for 6 hours. iPSC-ECs were stained with Calcein AM (green) before seeding. Pictures are under 10X magnification. In (C), (D), (H), (I), data shown as the mean ± SEM. *p<0.05, **p<0.01, control (n=4) vs HLHS (n=3). See also Figure S3.

Through binding to integrin receptors, different ECMs act as extracellular factors to initiate intracellular signaling cascades related to EndoMT, cell growth, apoptosis, and migration(Carey, 1991; Post et al., 2019; Schroeder et al., 2003; Short et al., 1998). Reduced integrin signaling predicted by scRNA-seq in iPSC-ECs was confirmed using an adhesion assay. Acute adhesion to several ECM components, such as collagen IV, gelatin, and laminin were profoundly suppressed in HLHS iPSC-ECs (Figure 3D). Intriguingly, cell proliferation was also reduced (Figure 3E), a possible sequela of reduced ECM-integrin interaction(Li et al., 2012). However, the altered integrin-ECM interactions did not affect cell migration in HLHS iPSC-ECs (Figure S3A).

In both myocardial and endocardial cells, NOTCH signaling plays a pivotal role during cardiac and valvular development(Luxan et al., 2016; MacGrogan et al., 2010). NOTCH mutation or haploinsuffciency are strongly implicated in congenital heart diseases incidence(Hrstka et al., 2017; Kobayashi et al., 2014; Theodoris et al., 2015; Yang et al., 2017). In our study, we also demonstrated that NOTCH signaling related genes were significantly down-regulated in HLHS iPSC-ECs compared with controls judged by scRNA-seq and qPCR analysis (Figure 3F-H). The imbalanced homeostasis of this process could cause abnormal valve formation during heart development.

Notch signaling can be activated by blood flow, which is an important stimulus for heart chamber formation(Andres-Delgado and Mercader, 2016). Under static condition, the expression levels of flow-responsive genes were reduced in iPSC-ECs from HLHS patients (Figure S3C). Interestingly, the differences between control and HLHS iPSC-ECs were diminished under 15dynes/cm^2^ laminar shear stress (LSS), as judged by cell alignment and gene expression levels (Figure S3B and S3C). This indicates that application of the appropriate blood flow may rescue HLHS iPSC-ECs shear response, even with an abnormal shear sensing system at basal level.

Vascular endothelial growth factor (VEGF) signaling, a strong pro-angiogenic factor, was suppressed in HLHS iPSC-ECs by ontology analysis (Figure 3A) and reduced VEGF expression has been described in the left ventricle from severe HLHS patients(Gaber et al., 2013). In accordance with the GO prediction, tube formation assays demonstrated a significant suppression of angiogenesis in HLHS iPSC-ECs compared with controls (Figure 3I).

### HLHS iPSC-ECs impeded cardiomyocyte proliferation and maturation

We hypothesized that a defective endocardium in HLHS could impair cardiomyocyte proliferation and contribute to the ventricular underdevelopment. To recapitulate the crosstalk between the endocardium and myocardium in human heart, we used a transwell co-culture system (Figure 4A) to culture normal iPSC-derived cardiomyocytes (iPSC-CMs) (Day 15-20) with normal or HLHS iPSC-ECs. RNA-seq analysis revealed distinct transcriptomic differences in cell cycle, cardiac conduction, cardiac muscle contraction, and sarcomere pathways when comparing iPSC-CMs co-cultured with control vs. HLHS iPSC-ECs (Figure 4B & C, Figure S4A). Similarly, KEGG pathway analysis also indicated repression of cardiac muscle contraction signaling (Figure S4B). These HLHS iPSC-EC-induced dysfunctional iPSC-CMs showed decreased proliferation with fewer Ki67^+^ cells (Figure 4D), as well as reduced gene expression levels of proliferation and maturation related markers (Figure 4E & F). These data predicts that the HLHS endocardium can hamper myocardial development by impairing valve formation, as well as cardiomyocyte growth and maturation.

**Figure 4.**
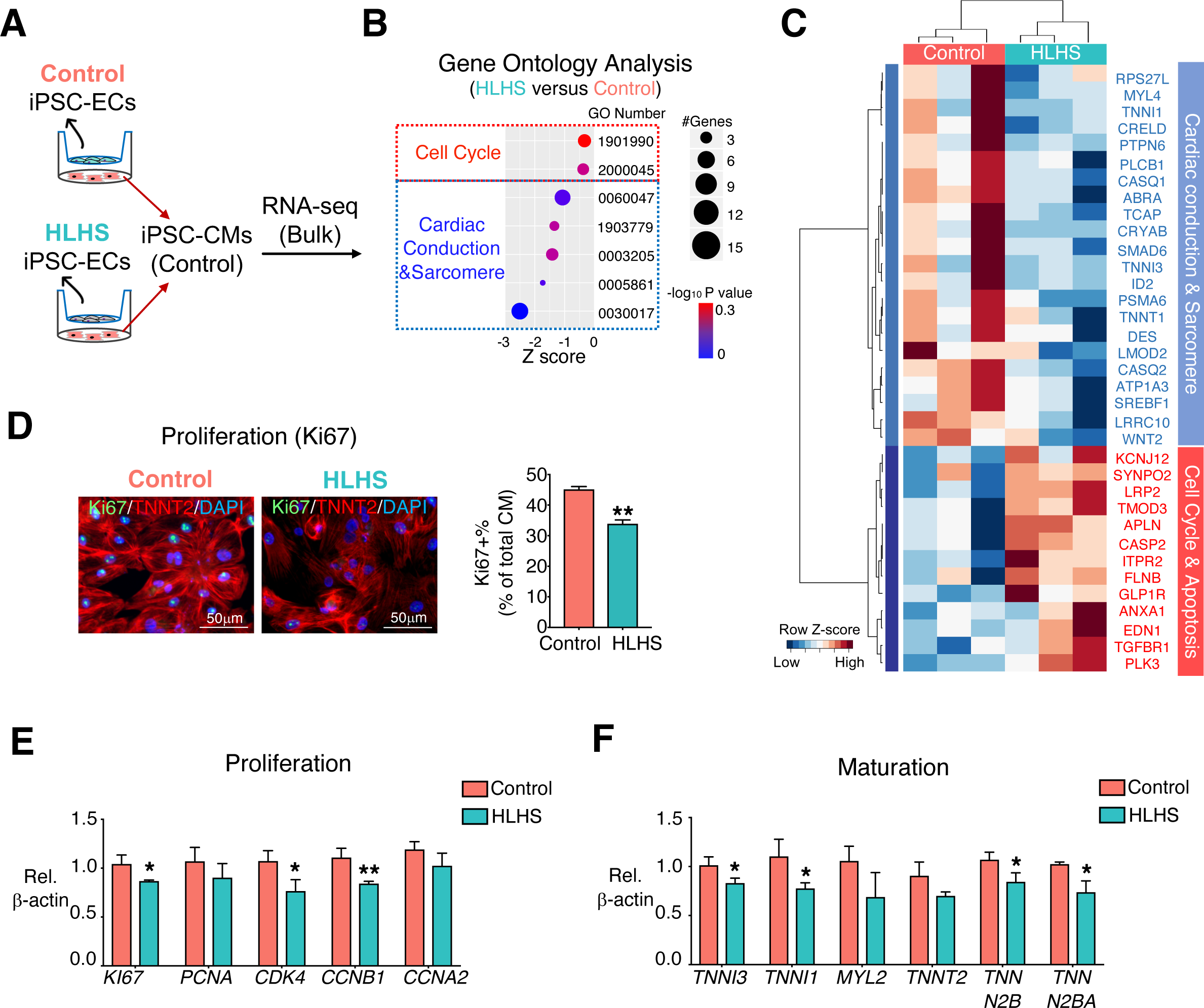
HLHS iPSC-ECs impeded cardiomyocyte proliferation and maturation. (A) Normal iPSC-CMs were co-cultured with iPSC-ECs from control or HLHS patients for 48 hours. (B) Transcriptomic profiling and functional enrichment analysis were performed on iPSC-CMs from (A) by bulk RNA-seq (n=3). −log10P indicates the significance of enrichment. Z-score defines the changing trend of enriched functions between HLHS vs control; z-score<0 means down-regulation in iPSC-CMs co-cocultured with HLHS iPSC-ECs. (C) Heatmap visualization of DEGs from GO terms in (B). (D) Immunostaining of Ki67 and TNNT2 in iPSC-CMs from (A). DAPI: nucleus; Green: Ki67 positive nucleus; Red: TNNT2 positive cardiomyocytes. (E) Gene expression related to proliferation and cardiac maturation in iPSC-CMs from (A). In (D)-(F), data shown as the mean ± SEM. *p<0.05, **p<0.01, control (n=3) vs HLHS (n=3). See also Figure S4.

### Endocardial homeostatic functions and endocardium-myocardium crosstalk are dependent on fibronectin 1, which is absent in HLHS

We next compared the differentially expressed genes (DEGs) in a normal left ventricle (LV) versus an underdeveloped left ventricle (ULV) from human fetal hearts (Figure S5A). The underdeveloped human fetal heart was micro-dissected, dissociated, and sequenced at single cell level. scRNA-seq analysis captured the major repertoire of the cardiac cell types (Figure S5B & C), including the endocardium (*NPR3^+^*), endothelium (*APLN^+^*), and cardiomyocytes (*TNNI3^+^*). Further analysis of CD144^+^ cells identified various EC populations (Figure S5D-G). Integration of scRNA-seq datasets from normal heart (day 83) and ULV (day 84) ECs showed similar endocardial and endothelial populations on the UMAP projection (Figure S5H-K). GO analysis of the DEGs between ULV versus control fetal heart showed dysregulated pathways in the ULV endocardial cell population (Figure S5L), similar to that observed in the HLHS iPSC-ECs scRNA-seq analysis (Figure 3A). These aberrant pathways include impaired focal adhesions, proliferation and increased cell death.

Genes altered in both HLHS iPSC-ECs and endocardial cells from ULV compared with controls were selected for further analysis (Figure 5A). In total, 259 DEGs (117 down-regulated and 142 up-regulated) were identified between ULV vs. normal human fetal left heart endocardium, while 216 DEGs (190 down-regulated and 26 up-regulated) were found in HLHS vs normal iPSC-endocardial cells (Figure S6A). Of those, 16 DEGs were shared by both endocardial cells isolated from the human fetal heart with ULV and HLHS iPSC-endocardial cells (Figure S6B). Fibronectin 1 (*FN1*) was one of the DEGs significantly repressed in both groups (Figure 5A). The suppression of *FN1* from scRNA-seq was confirmed by qPCR in 3 HLHS iPSC-EC lines compared with 4 control lines (Figure 5B). In normal human fetal heart, FN1 was universally expressed but enriched in coronary ECs (Figure S6C, left panel), ventricular endocardium (Figure S6B, middle panel), and atrial endocardium (Figure S6B, right panel). Consistent with the reduction at transcript level, FN1 protein expression was significantly reduced in the endocardial layer of HLHS patient hearts compared with control hearts (Figure 5C). FN1 promotes cardiomyocyte proliferation(Hornberger et al., 2000; Ieda et al., 2009), and is required for normal heart development(Mittal et al., 2013), as well as for the left-right asymmetry in vertebrates(Pulina et al., 2011). Therefore, we speculated that FN1 suppression may partially contribute to the ventricular underdevelopment in at least a subset of HLHS cases through the modulation of endocardial cell functions.

**Figure 5.**
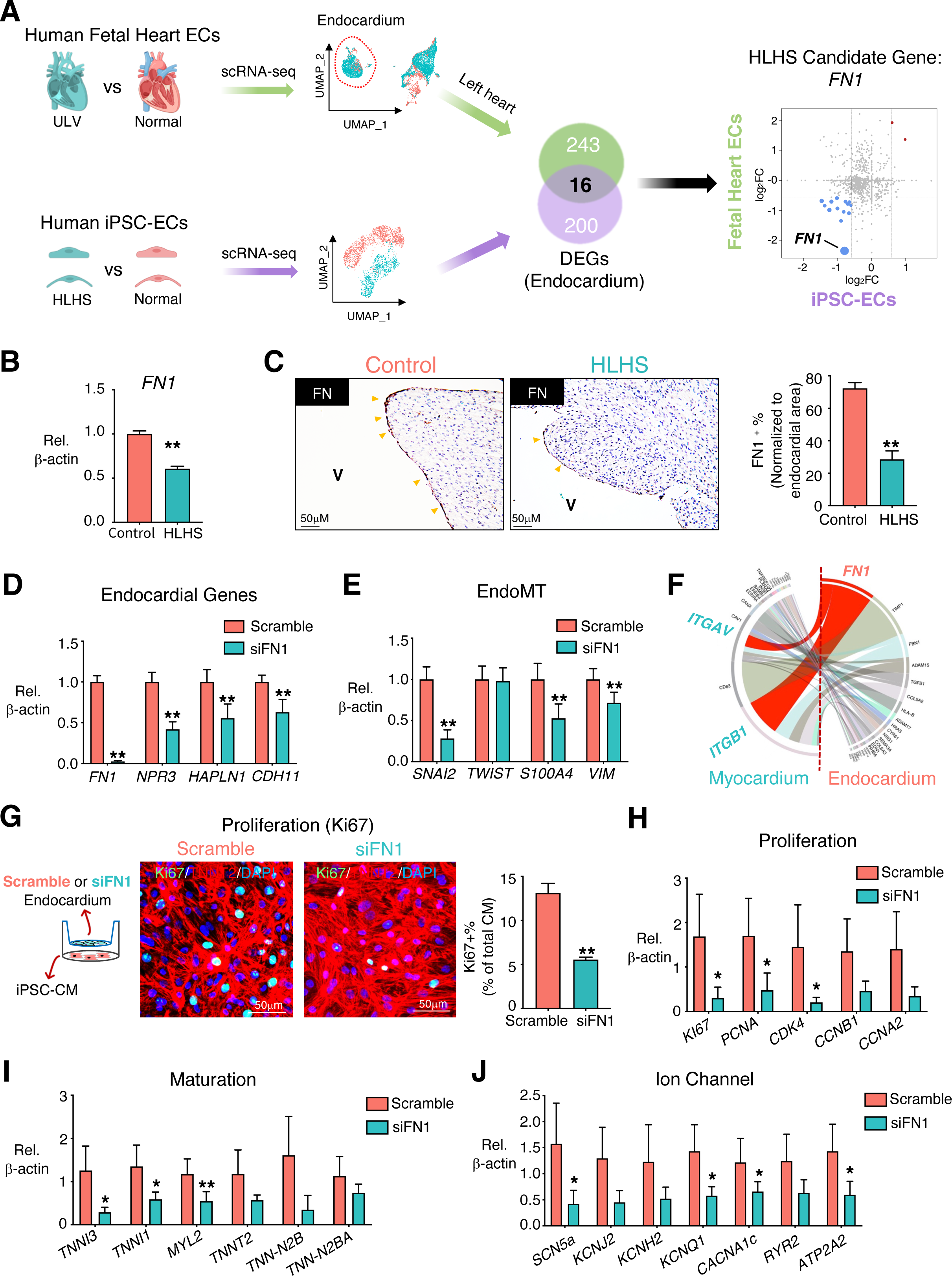
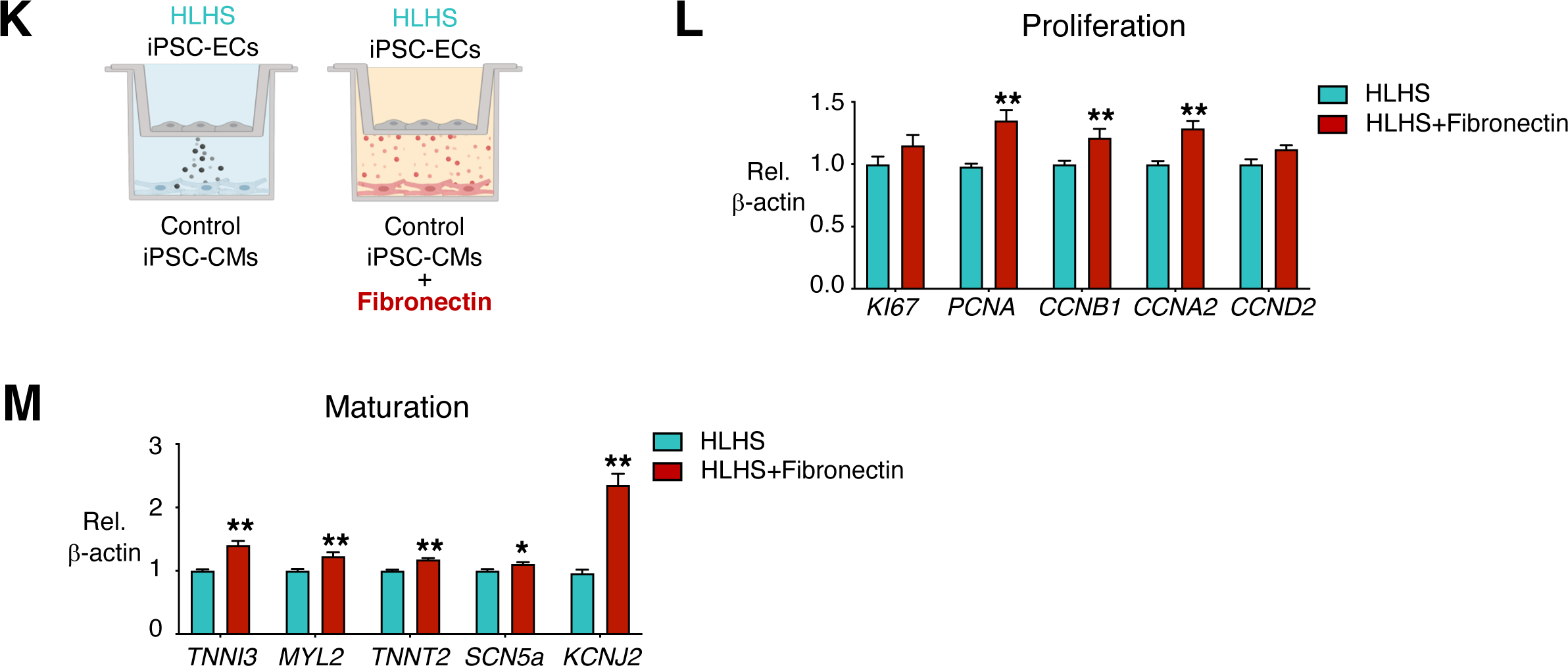
Endocardial homeostatic functions and endocardium-cardiomyocyte crosstalk is dependent on FN1, which is absent in HLHS. (A) Illustration: Overlap of the DEGs from human fetal heart tissue and iPSC-ECs comparing control vs. patient. DEGs from human fetal heart tissue were determined based on the transcriptomic comparison of the endocardial subpopulation from healthy control vs. fetal heart with underdeveloped left ventricle using scRNA-seq analysis. Monte-Carlo simulation indicated the overlapping significance. (B) *FN1* gene expression in iPSC-ECs from control (n=4) and HLHS patients (n=3). (C) Left panel: Immunostaining of FN1 proteins at endocardial layers in human hearts from control and HLHS patients. Yellow arrowheads indicated positive FN1 staining. V: ventricular chamber. Right panel: Quantification of FN1 positive endocardial cells in control (n=6) and HLHS (n=12) patients. Expression of genes related to the endocardium (D) and EndoMT pathways (E) in left heart endocardial cells with *FN1* knock-down. (F) The interactions strength of each endocardium ligand and its cardiomyocyte receptor are displayed in the chord diagram based on day83 normal human fetal heart scRNA-seq. A ligand-receptor relationship is marked by the color of the ligand (FN1 in red), and interaction strength is shown by the width of chords. Left half circle: Ligands expressed in endocardium; right half circle: receptors expressed in myocardium. (G) Immunofluorescence staining of Ki67 protein in normal iPSC-CMs after co-cultured with primary endocardium with or without *FN1* knock-down. DAPI: nucleus; Green: Ki67 positive nucleus; Red: TNNT2 positive cardiomyocytes. Gene expression related to proliferation (H) and cardiac maturation (I) and ion channels (J) in iPSC-CMs from (G). (K) Illustration of iPSC-CMs and iPSC-ECs co-culture experiments with supplementation of fibronectin (5μg/ml, 48 hours) to the iPSC-CMs. qPCR quantification of gene expressions related to proliferation (L) and myocardial maturation (M). (D-M), n=3. Data shown as the mean ± SEM. *p<0.05, **p<0.01, control vs HLHS, scramble vs siFN1, or HLHS vs HLHS + fibronectin. See also Figures S5 and S6.

As iPSC-ECs are a mixed population with both endocardial and endothelial cells, we established a protocol to purify each of these two populations directly from human fetal heart to test the endocardial-specific effect of FN1. The innermost layer of left or right ventricular free wall was immersed into the digestion buffer for several minutes, and then the detached endocardial cells were collected for CD144 antibody selection. Similarly, the outermost ventricular free walls and aorta were digested and further purified to generate the EC population (Figure S6D). Endothelial gene expression assessment revealed enriched EC populations as compared to non-EC populations such as fibroblasts and cardiomyocyte (Figure S6E). Moreover, enriched endocardial and endothelial populations can be achieved using our isolation and purification protocol: endocardial markers, *NPR3* and *HAPLN1* were selectively expressed in human heart endocardial cells when compared to coronary ECs or aortic ECs; whereas coronary markers, *MGLL* and *APLN* showed higher expression in the isolated coronary ECs (Figure S6F).

Next, we reduced *FN1* expression level using siRNA in the native endocardial cells isolated from fetal hearts and found significantly decreased endocardial (Figure 5D) and EndoMT genes (Figure 5E), similar to those observed in HLHS iPSC-ECs (Figure 2 & 3). scRNA-seq from normal fetal heart predicted FN1-interacting partners in the cardiomyocyte population, *ITGA5* and *ITGA1* (Figure 5F). Suppression of *FN1* in the endocardium not only impaired its intrinsic functions, but also significantly impeded normal iPSC-CMs functions, i.e. there were fewer proliferating CMs judged by reduced Ki67^+^ cells (Figure 5G) and reduced expression of genes related to proliferation (Figure 5H). Additionally, the iPSC-CMs co-cultured with endocardial cells with reduced FN1 had decreased expression of genes related to maturation and ion channels (Figure 5I & J). Intriguingly, addition of fibronectin to the control iPSC-CMs co-cultured with HLHS iPSC-ECs significantly improved CMs proliferation and maturation (Figure 5K-M). These data suggest that reduced FN1 in HLHS endocardial cells may underlie the impaired myocardial development in HLHS, which was consistent with the phenotype in iPSC-CMs co-cultured with HLHS iPSC-ECs (Figure 4).

### HLHS *de novo* mutations altered endocardial and myocardial functions through transcriptional regulation of endocardial genes

As revealed by the fetal heart scRNA-seq, most of the genes that have been reported to harbor DNMs in HLHS patients were highly expressed in the endocardium/endothelium (Figure 1D). The expression levels of several of these genes in HLHS iPSC-ECs were consistently down-regulated compared with controls, such as *ETS1, CHD7, KMT2D, ARID1B,* and *FOXM1* (Figure 6A and Figure S7A), and these are primarily chromatin remodelers and transcription factors (Figure 6A, gene names highlighted in red). Damaging DNMs accounting for ∼10% of the sporadic CHD cases are highly enriched in genes encoding chromatin remodelers(Jin et al., 2017). Reducing levels of *ETS1*, *CHD7*, and *KMT2D*, but not *ARID1B* or *FOXM1* (Figure S7B-D) in human primary endocardial cells led to the suppression of endocardial-related genes (Figure 6B) and *FN1* expression (Figure 6C). To further understand the role of these mutated genes in regulating downstream endocardial gene network, we first focused on one transcription factor *ETS1* and one chromatin remodeler *CHD7*. Chromatin immunoprecipitation assays (ChIP) on iPSC-ECs found reduced ETS1 and CHD7 binding to the *NPR3* promoter region in HLHS iPSC-ECs compared to the control (Figure 6D). We also applied HUVEC tracks from ENCODE to detect a region in the FN promoter (‘FN1-a’ in Figure 6E) and enhancer (‘FN1-b’ in Figure 6E) that we used for the ChIP-qPCR. Both ETS1 and CHD7 showed reduced binding to *FN1* promoter and enhancer in HLHS vs. control iPSC-ECs (Figure 6E). Similarly, their binding capacities to the enhancer regions (i.e., H3K27ac and H3K4me high, while H3K4me3 low) of another pivotal endocardial gene, *CDH11* were consistently reduced (Figure S7E).

**Figure 6.**
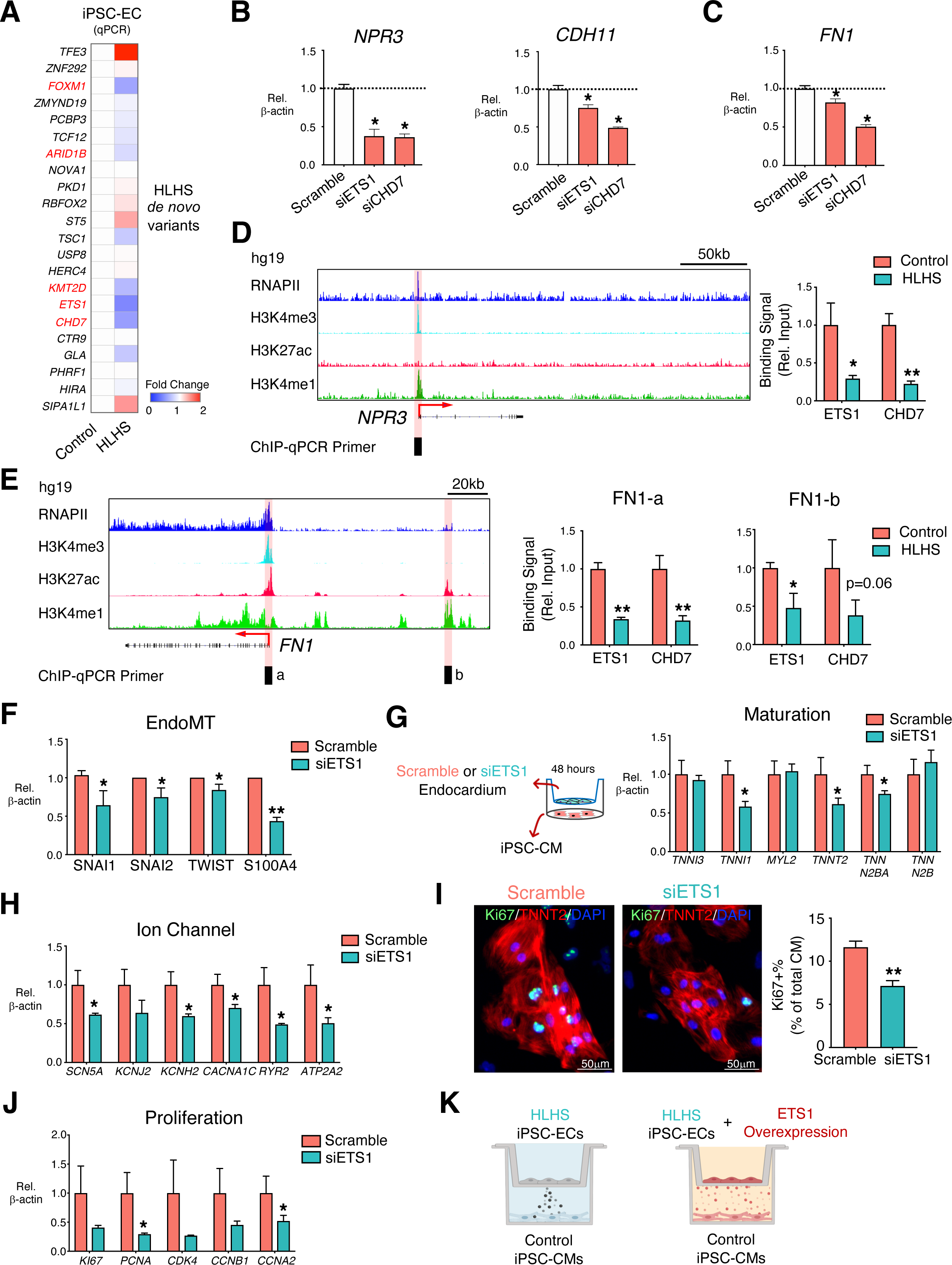
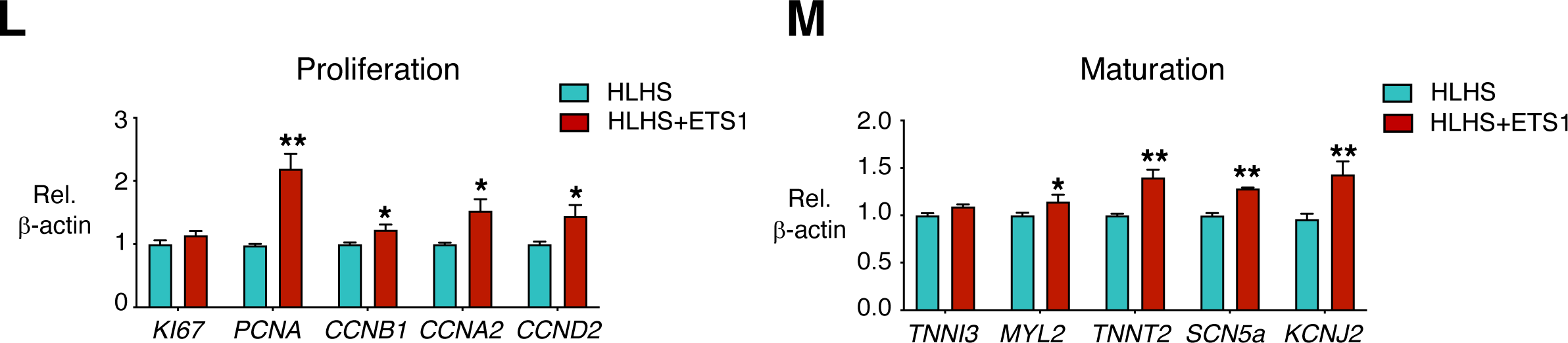
HLHS DNMs altered endocardial and myocardial functions through transcriptional regulation of endocardial genes and FN1. (A) qPCR detection of HLHS DNM genes in iPSC-ECs from control (n=4) and HLHS patients (n=3). Red denotes high expression, blue minimal expression. Genes not shown mean non-detectable in iPSC-ECs. Genes highlighted by red are transcription factors or chromatin modifiers. qPCR validation of the endocardial gene expression (B) and *FN1* (C) in primary human fetal endocardial cells after *ETS1* or *CHD7* knock-down, respectively. (D) Left: RNAPII and histone marker ChIP-seq tracks of the *NPR3* gene in HUVECs from the ENCODE. The black box and red shade indicate the designated location of primers for ChIP-qPCR. The red arrow indicates the transcriptional start site (TSS) and direction of transcription. Right: ChIP-qPCR of ETS1 or CHD7 binding to the *NPR3* promoter region in control and HLHS iPSC-ECs. (E) Left: Histone ChIP-seq tracks of the *FN1* gene in HUVECs from the ENCODE. Right: ChIP-qPCR for ETS1 or CHD7 to the *FN1* promoter (a) and enhancer (b) regions. *p<0.05, **p<0.01, control (n=3) vs HLHS (n=3). Data are represented as mean ± SEM. (F) EndoMT-related gene expression in endocardial cells after knock-down of *ETS1*. (G) iPSC-CMs were co-cultured with isolated fetal left heart endocardial cells with or without *ETS1* knock-down. Expression of genes related to cardiomyocyte maturation (G) and ion channel (H) in iPSC-CMs were examined by qPCR. (I) Immunostaining of Ki67 (green) and TNNT2 (red) in iPSC-CMs from (G). DAPI: nucleus. (J) qPCR for proliferative genes in iPSC-CMs from (G). (K) Illustration of iPSC-CMs and iPSC-ECs co-culture experiments with ETS1 overexpression in HLHS iPSC-ECs (HLHS+ETS1). qPCR quantification of gene expressions related to proliferation (L) and cardiomyocyte maturation (M). n=3 patients in each treatment group. Data shown as the mean ± SEM. *p<0.05, **p<0.01, scramble vs. siRNA knockdown or HLHS vs. HLHS+ETS1. See also Figure S7.

To further understand how the genes associated with HLHS regulate downstream endocardial and myocardial dysfunctions, we first focused on *ETS1* since it was the utmost down-regulated gene of the group (Figure 6A). *ETS1* expression was also reduced in HLHS iPSC-ECs based on scRNA-Seq analysis (Figure S7F). Reduction of *ETS1* expression using siRNA in primary human endocardial cells not only significantly suppressed genes related to the endocardium and FN1 (Figure 6B & C), but also reduced EndoMT signaling (Figure 6F). Mechanistically, knockdown of *ETS1* in human endocardial cells diminished RNAPII recruitment to the promoter regions of endocardial genes, such as *NPR3*, *PLVAP*, and *CDH11* but not *FN1* (Figure S7G). We speculated that other chromatin modifiers, such as mediator complex or the dynamic RNAPII regulatory steps during transcription initiation might be involved in ETS1-mediated *FN1* transcription(Chen et al., 2015; Jeronimo and Robert, 2017; Steurer et al., 2018). Correspondingly, we analyzed a previously published dataset of *ETS1* over-expression (Mod_ETS1 vs Mod_GFP) in human umbilical vein endothelial cells (HUVECs)(Chen et al., 2017). RNAPII binding capacity to endocardial gene promoter regions were significantly increased with *ETS1* over-expression. For example, the promoter regions of the endocardial genes, *NPR3* and *PLVAP* (ENCODE HUVECs datasets), were enriched with RNAPII signaling under *ETS1* overexpression (Mod_ETS1 vs Mod_GFP) (Figure S7H), which was in line with our *ETS1* knock-down panel. Additionally, similar to the HLHS iPSC-ECs and the *FN1*-deficit endocardium, endocardial cells lacking *ETS1* impeded CM maturation (Figure 6G & H) and proliferation, evidenced by reduced Ki67^+^ CMs (Figure 6I) and less proliferative gene expression levels (Figure 6J). Intriguingly, iPSC-CMs co-cultured with HLHS iPSC-ECs over-expressing *ETS1* increased the proliferation and maturation of the CMs (Figure 6K-M). In conclusion, these DNMs provide a genetic clue to explain the mechanism involved in the endocardial defects and reduced *FN1* in HLHS.

### Suppression of ETS1 in *Xenopus* caused reduced endocardial FN1 and impaired heart development

Previously it has been demonstrated that loss of Ets1 in the cardiac mesoderm of *Xenopus* embryos leads to heart malformations exhibited by reduced ventricular chamber size, similar to that of a HLHS heart (Nie and Bronner, 2015). To determine whether the loss of Ets1 results in dysregulated FN1 expression in *Xenopus* heart, similar to the results above from our *in vitro* and human studies, we examined the expression of FN1 by immunostaining. Ets1 was knocked-down in *Xenopus* hearts by injecting morpholinos (Ets1-MO) into cardiogenic mesoderm during cleavage stages. The suppression of Ets1 resulted in significantly reduced ventricular chamber size (outlined with a white dashed box) when compared to the controls (Figure 7A & B). In the control hearts, FN was detected in the inner lining of the endocardial tissue (n=4/4). In contrast, FN staining was lost in the endocardium of Ets1 morphant hearts (n=6/6) (Figure 7A & C). This *in vivo* evidence suggests the existence of an ETS1-FN1 regulatory axis that is sufficient to cause heart ventricular underdevelopment, similar to what is observed in HLHS.

**Figure 7.**
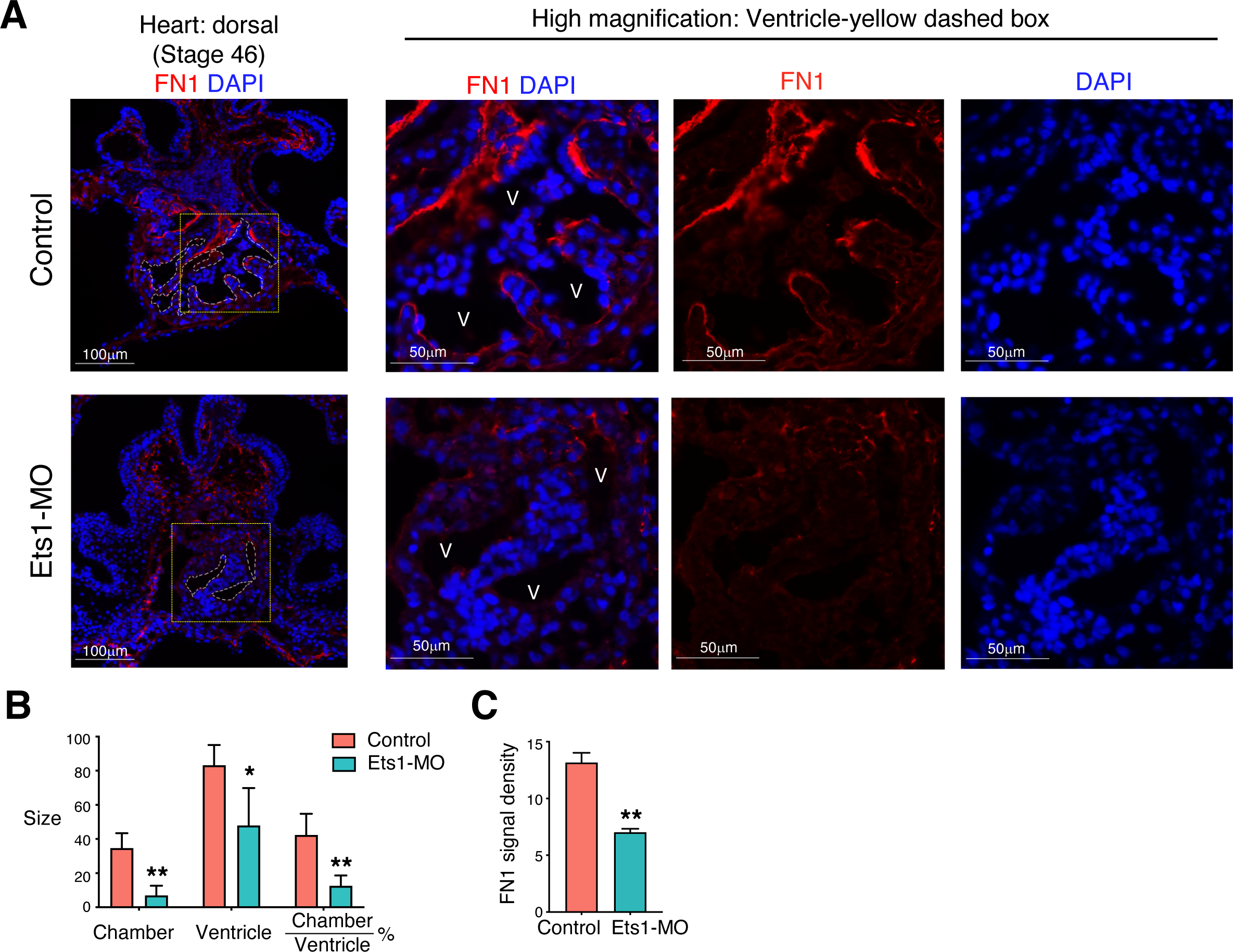
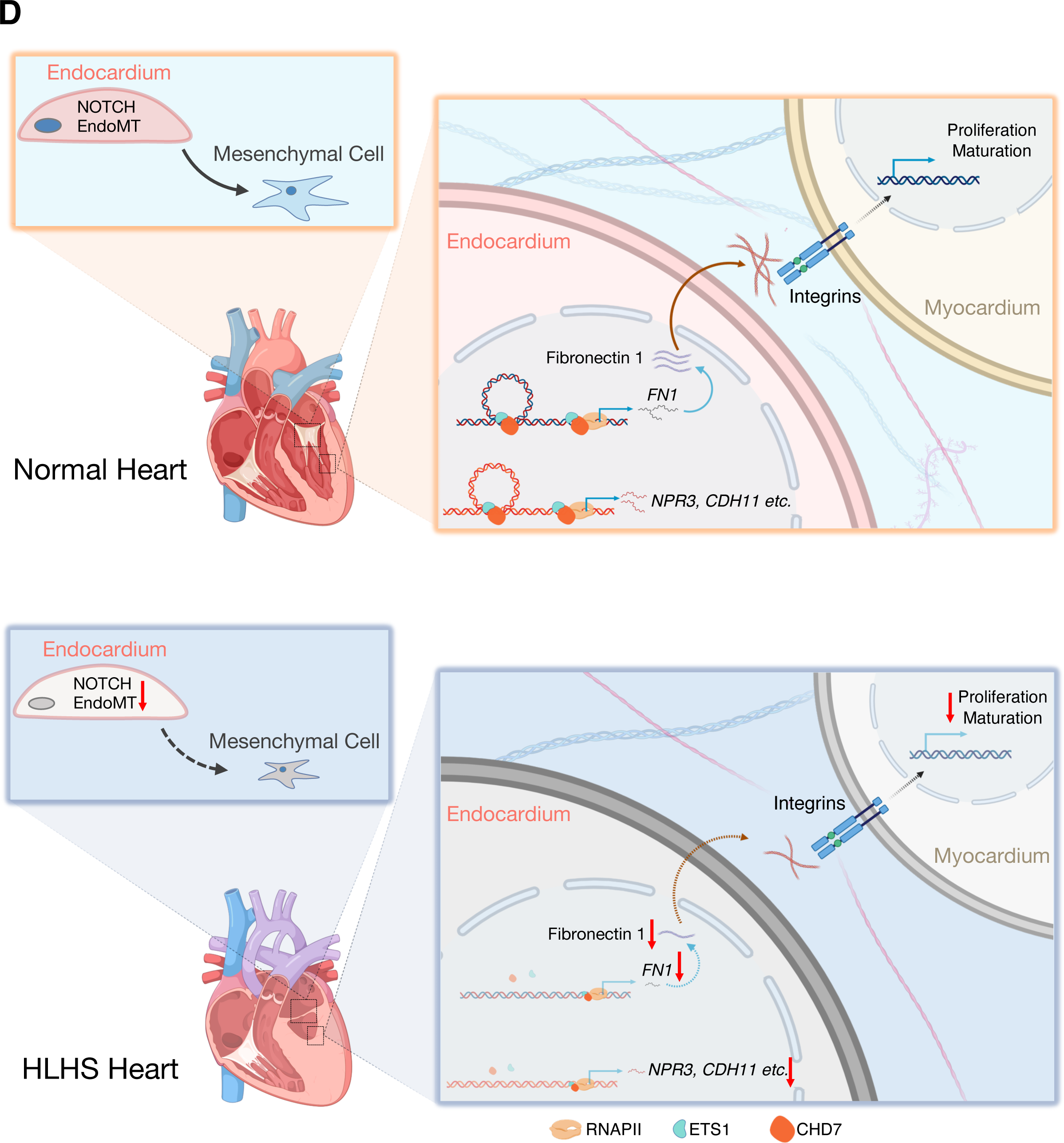
Knock-down of ETS1 *in vivo* caused FN1 reduction in the endocardium and leads to abnormal heart development. (A) Staining of FN1 in *Xenopus* heart with control or Ets1-MO injection. White dashed lines outline the chamber of the *Xenopus* ventricle. Yellow dashed box area was further zoomed in to visualize endocardial FN1 expression. V: ventricle. (B) Quantification of the absolute sizes of the heart chamber, ventricle, and the relative chamber size. (C) FN1 fluorescence signal intensity within the endocardium. control=4; Ets1-MO=6. Data shown as the mean ± SEM. *p<0.05, **p<0.01, control vs Ets1-MO. (D) Schematic illustration of the endocardial and myocardial defects in HLHS. In HLHS endocardial cells, ETS1 and CHD7 expression levels were significantly reduced compared to healthy controls. This directly led to the decreased expression and secretion of fibronectin, which further disrupted the fibronectin-integrin interaction with cardiomyocytes. Impaired integrin activation in cardiomyocytes resulted in decreased cell maturation and proliferation, which may contribute to the underdevelopment of the left ventricle. Additionally, decreased ETS1 and CHD7 also led to the suppression of downstream endocardial gene expression such as *NPR3* and *CDH11*. The impaired endocardium showed aberrant endothelial to mesenchymal transition as well as NOTCH signaling pathway, which are critical for cardiac valve development.

## Discussion

Myocardial abnormalities have been considered to be the main cause underlying hypoplasia of the left ventricle. Several studies have demonstrated impaired cardiogenesis and/or augmented myocardial apoptosis in iPSC-CMs or heart tissues from HLHS patients and animal models(Gaber et al., 2013; Hrstka et al., 2017; Kobayashi et al., 2014; Liu et al., 2017). However, growing evidence suggests that underdevelopment of the left ventricle could also due to the abnormal blood flow, which is secondary to hypoplastic and/or atretic valve structures. During heart development, the endocardium serves as a key component in facilitating valve, septum, and coronary artery formation and maturation. Despite its pivotal role, the potential mechanisms of endocardial defects in causing cardiac malformation in HLHS are unknown. Previously, it has been reported that genetic ablation of the endocardial cell population during murine cardiac development arrested the growth of the left ventricle, resulting in embryonic lethality by E11(Snider et al., 2014). Clinically, in the anatomic subset of mitral stenosis and aortic atresia, there is often a thickened layer of fibroelastosis lining the cavity of the ventricle in HLHS, indicative of endocardial injury(Grossfeld et al., 2019). Herein, using patient-specific iPSCs, fetal heart tissue with an underdeveloped left ventricle, and scRNA-Seq, we have provided the first direct evidence that severely impaired endocardial cell populations are present in HLHS patient hearts, and that these cell population alterations lead to an imbalanced functional homeostasis involving in the pathogenesis of HLHS.

The genetics underlying HLHS are complex. To better understand the mutations associated with CHD, the US National Heart, Lung, and Blood Institute (NHLBI) Pediatric Cardiac Genomics Consortium (PCGC) carried out whole exome sequencing (WES) in over 10,000 CHD probands. They showed that approximately 10% of the cases were attributable to DNMs, including significant enrichment in chromatin modifiers, which may have significant impact on downstream gene expression networks(Homsy et al., 2015; Zaidi et al., 2013). Specifically, there were 27 DNMs associated with HLHS phenotypes. Surprisingly, based on our single cell heart atlas, we found that 25 out of the 27 genes that harbored these DNMs were highly expressed in the endocardial and endothelial cell populations during heart development. This indicated that endocardial defects could play a major role in causing HLHS.

Interestingly, one HLHS patient carried only one of these DNMs. We hypothesized that there may be a common gene or pathway downstream of the genes harboring these DNMs that leads to the development of HLHS. Our study first focused on two endocardial genes whose expressions were significantly decreased in HLHS iPSC-ECs compared with controls: a transcription factor (*ETS1*) and a chromatin remodeler (*CHD7*). A loss-of-function mutation in *ETS1* has been identified in patients with hypoplastic left ventricle and other features that are seen in Jacobsen syndrome(Glessner et al., 2014; Tootleman et al., 2019; Ye et al., 2010). Genetic ablation of *ETS1* specifically in the cardiac mesoderm in *Xenopus* produced a HLHS like phenotype associated with loss of endocardial cells (Nie and Bronner, 2015). Likewise, systemic deletion of *ETS1* in mice caused a non-apex forming left ventricle phenotype(Ye et al., 2010), one of the hallmarks of HLHS. However, the penetrance of the *ETS1* mutation is low, indicating that there might be several other rare or even common variants that are responsible for the HLHS phenotypes. A loss-of-function mutation in *CHD7* was related to severe congenital heart defects in CHARGE syndrome, with overrepresentation of atrioventricular septal defects and outflow tract defects(Corsten-Janssen and Scambler, 2017). Because there is a large gene network downstream of *ETS1* and *CHD7*, we first narrowed down the pool of candidate genes by focusing on the ones that were differentially expressed between control vs. HLHS endocardial cells. *FN1* was the most strikingly down-regulated gene in HLHS endocardium. Interestingly, through ChIP-qPCR analysis, we also found that the binding of both ETS1 and CHD7 to the promoter and enhancer regions of *FN1* was significantly decreased in HLHS iPSC-ECs compared with controls.

FN1 is one of the major extracellular matrix (ECM) components that plays active roles in cardiac development. It has been reported that proliferating human fetal CMs synthesized endogenous fibronectin, while inhibition of its production arrested CM growth(Hornberger et al., 2000). Furthermore, exogenously coating a culture dish with fibronectin augmented embryonic CM proliferation(Ieda et al., 2009). *In vivo*, FN1-null mice exhibited abnormal left ventricle development at the embryonic stage, possibly resulting from aberrant differentiation of cardiac precursors(Mittal et al., 2013). Therefore, FN1 may act as an extracellular factor to trigger a cascade of signaling pathways that modulate CM growth. There is accumulating evidence that supports integrin *α*5*β*1 as the key receptor involved in FN1 signaling in CMs (Ieda et al., 2009; Mittal et al., 2013). We further confirmed this in our ligand-receptor interaction analysis using our human fetal heart scRNA-seq data. Herein, we provided that the endocardium as another important source of FN1 to facilitate normal heart development; when FN1 is lacking, it can lead to impaired iPSC-CM growth through disruption of the FN1-integrin *α*5*β*1 interaction. When FN1 is supplemented to the HLHS iPSC-ECs and iPSC-CMs co-culture system, it rescues the decreased proliferation and maturation of the CMs.

Deep understanding of the endocardial defect in HLHS patients requires an effective protocol to differentiate patient-specific iPSCs into endocardial cells. However, to date, there is no such protocol published. The most relevant cardiac EC differentiation protocol was established by Palpant et al. (Palpant et al., 2017), which could distinguish cardiac ECs and hemogenic ECs. Despite this caveat, we were able to utilize scRNA-seq to assess the endocardial specific gene expression abnormalities in HLHS by separating different EC subpopulations from patient-specific iPSCs differentiated into the heterogenous iPSC-ECs.

The first scRNA-seq work that delineated the human heart developmental track identified four different cardiac EC populations, including free wall endocardium, coronary vascular ECs, vascular ECs, and valvar ECs(Cui et al., 2019). However, due to the low numbers captured in their EC population, it would be difficult to perform in-depth EC sub-cluster analysis with the given dataset depth. In our current study, we specifically enriched cardiac EC populations and established a high resolution human cardiac EC atlas including coronary artery ECs, vein ECs, capillary ECs, endocardial ECs, valvular ECs, and lymphatic ECs. Our study provides a more comprehensive database to study the differences between these EC subtypes in human, and can be applied to study other EC-related congenital heart diseases.

Collectively, our studies based on scRNA-seq analysis using patient-specific iPSC-ECs and human fetal heart tissue reveal robust endocardial defects and aberrant endocardium-myocardium crosstalk in HLHS. By modulating HLHS related gene variants and ChIP-qPCR analyses, we further elucidated molecular mechanisms underlying the dysfunctional endocardium (Figure 7D). Our study provides a new platform to improve our understanding of HLHS etiology in a patient-specific manner, and paves the way to develop novel therapies for this condition in the future.

## Acknowledgments

We thank Drs. Jingjing Li, Qing Liu, Michael P. Snyder and Mr. Eric Wei for DNM analysis and scRNA-seq sample preparation. We thank Dr. Virginia D. Winn for helping us with human fetal tissue collection. We greatly appreciate the administrative help of Ms. Michelle Fox. We also thank Drs. Mark Mercola, Kristy Red-Horse, James Wells, and Kyle Loh for providing intellectual consultant to the project. We also thank ReGen Theranostics, Inc Rochester, MN as the manufacturer for the iPSC lines and Dr. William Pu from Boston Children’s Hospital, Harvard Medical School who provided GFP and ETS1 overexpression lentivirus. This work was supported by single ventricle gift fund from Stanford University, Todd and Karen Wanek Family Program for Hypoplastic Left Heart Syndrome, NIH 2R24HD000836-52 from the Eunice Kennedy Shriver National Institute of Child Health and Human Development (The University of Washington Birth Defects Research Laboratory), NIH 5T32HL098049-10 (MM), NIH R01 HL145675, NIH R01 HL141371 (JCW), and NIH K99 HL135258 (MG).

## Author Contributions

YM designed and performed experiments, analyzed data and wrote the manuscript. LT designed and analyzed the scRNA-seq and Bulk RNA-seq data and wrote the manuscript. MM analyzed the imaging data and revised the manuscript. SP and FG provided iPSC-CMs and helped with experimental designs. JL and SN performed Ets1 knock-out in *xenopus* heart and analyzed the results. AG helped with knock-down and qPCR assays. YW performed Ligand-receptor interaction analysis and consulted on scRNA-seq analysis. JM performed and analyzed shear-stress assay. HZ and NM contributed to iPSC line maintenance, cell sorting, and provided human cardiac fibroblasts. BZ offered substantial intellectual contribution to ETS1 ChIP-seq analysis. SM and DC provided human heart tissue sections and revised the manuscript. TN provided HLHS iPSC lines. JW, MR, and SW offered substantial intellectual contribution to the project and revised the manuscript. MG designed the studies, oversaw the scRNA-seq, EC functional assays, data acquisition and analysis, and manuscript preparation and editing.

## EXPERIMENTAL MODEL AND SUBJECT DETAILS

### Human cell lines

Clinicopathological baseline characteristics of patients and donors are given in Table 1. De-identified HLHS patients’ peripheral blood mononuclear cells (PBMCs) were collected at Mayo Clinic through the Todd and Karen Wanek Family Program for Hypoplastic Left Heart Syndrome. HLHS iPSC lines 1, 2, and 3 were reprogrammed by ReGen Theranostics, Inc Rochester, MN. Skin biopsies from healthy control individuals (control cell line 1, 3, and 4) were obtained from Stanford Transplant Center, and skin fibroblasts were further reprogrammed at Stanford University Cardiovascular Institute (SCVI) Biobank. Control cell line 2 was purchased from CIRM iPSC Repository (Cat# CW60359EE1). The derivation protocol was modified based on previous publications (Burridge et al., 2014; Gu et al., 2017). Consent was obtained from both control and patients under approved IRBs: Mayo Clinic: 10-006845; Stanford: IRB 5443.

### Human heart tissue sections

Samples from the heart free wall were acquired from the pathology department at the time of fetal autopsies performed as soon as possible after pregnancy termination or fetal demise. All investigations were conducted according to the Declaration of Helsinki principles. Studies were approved by the Hospital for Sick Children and the Mount Sinai Hospital institutional review boards, and written informed consent was obtained from study participants (pregnant mothers) before inclusion in the study. Myocardial tissues were fixed and sectioned (5 μm) at University of Toronto and sent to Stanford University for staining.

### Frog embryo

The *Xenopus laevis* embryos were obtained by *in vitro* fertilization from wild type adult frogs (Nasco). The sex of embryos was not determined. All experimental procedures involving frog embryos were performed according to USDA Animal Welfare Act Regulations and have been approved by Institutional Animal Care and Use Committee, in compliance of Public Health Service Policy.

## METHOD DETAILS

### Cell culture

#### Endothelial differentiation

For differentiation of endocardial/endothelial cells from iPSCs, we used our previously published protocol(Gu, 2018; Gu et al., 2017). Briefly, iPSCs at 80% confluence were placed in differentiation medium (RPMI and B27 supplement minus insulin, Gibco) with 6 μM CHIR-99021 (Selleck Chemicals) for two days, followed by 3μM CHIR-99021 for another two days. Then medium was switched to EGM-2 EC differentiation medium (Lonza) with 50 ng/mL vascular endothelial growth factor (VEGF, Gemini) and 10 ng/mL basic fibroblast growth factor (bFGF, Gemini) for 5 days. On the last day of differentiation, cells were dissociated and sorted for CD144^+^ population using a MACS sorter (Miltenyi Biotec). After sorting, the cells were plated on 0.2% gelatin (Sigma)-coated plate with EGM-2 EC culture medium until 80%-90% confluence. iPSC-ECs used in the current study were between passage 2-4.

For differentiation of cardiomyocytes, we used previous published protocol (Burridge et al., 2014) iPSCs were maintained in E8 media (Gibco). At a confluency of 70-75%, iPSCs were passaged at a 1:12 ratio with Accutase (Gibco) or 0.5mM EDTA (Gibco) and evenly plated on 6-well plates. At a confluency of 95-98% on Day 0, E8 media was changed to RPMI 1640 (Gibco) containing 6 µM CHIR-99021 (Selleck Chemicals) and B27 Minus Insulin (Gibco). On Day 2, media was changed to RPMI 1640 supplemented with B27 Minus Insulin. On Day 3, media was changed to RPMI containing 2 μM C59 (Selleck Chemicals) and B27 Minus Insulin. On day 5, cells were rinsed with 1x PBS, and fed with RPMI 1640 supplemented with B27 Minus Insulin. On day 7, media was changed to RPMI 1640 supplemented with B27 Plus Insulin (Gibco). By day 9, beating cells could be observed and cells were glucose starved to enrich for cardiomyocytes by two-day treatment with RPMI 1640 Minus Glucose supplemented with B27 Plus Insulin. On day 12-13, cardiomyocytes were replated using 10X TRYPLE (Gibco) Select for 10 minutes. Cells were detached from wells by gentle pipetting and neutralization of TRYPLE Select by cardiomyocyte replating media consisting of 80% RPMI 1640, 20% Knockout Serum (Gibco), B27 Plus Insulin, and 10 µM Y-27632 (Tocris). The day after replating, cardiomyocytes were subsequently maintained in RPMI 1640 with B27 Plus Insulin by media changes every two days.

#### Frog embryo manipulations

*Xenopus laevis* embryos were obtained and microinjected with Ets1-MO as previously described (Nie and Bronner, 2015). Ets1-MO (5’-TAAGGTCTAGTGCAGCTTTCATGGC-3’) was injected into both dorsal vegetal blastomeres at 8-cell stage to target cardiac mesodermal tissue. All experimental procedures were performed according to USDA Animal Welfare Act Regulations and have been approved by Institutional Animal Care and Use Committee, in compliance of Public Health Service Policy.

#### Chromatin immunoprecipitation

ChIP assays were performed as previously described(Miao et al., 2018). Briefly, 1×10^7^ human fetal endocardial cells or iPSC-ECs were treated with 0.75% formaldehyde for 20 minutes at room temperature. Afterwards, fixation was stopped by adding 125mM glycine and cells were collected. The pelleted cells were lysed and sonicated using default parameter by Bioruptor® Pico (Diagenode Inc.) for 10 times using ‘30 seconds ON, 30 seconds OFF’ program at 4 °C. The sonicated samples were then centrifuged and 1% of supernatant was taken as input. After sonication, the chromatin was immunoprecipitated by various antibodies (ETS1, #39580, Active Motif-4μg per ChIP; RNA Pol II, #39097, Active Motif-4μg per ChIP; CHD7, #6505S, Cell Signaling Technology-4μg per ChIP) conjugated to pre-washed Protein A or Protein G Dynabeads. Protein and RNA were digested by proteinase K and RNase A, respectively. The purified chromatin DNA was then used as the template for a quantitative polymerase chain reaction. As an isotype control, non-specific IgG derived from the same species as specific antibodies were used in ChIP.

#### Quantitative Reverse-Transcription PCR

Our detailed protocol was previously published (Gu et al., 2017). Briefly, total RNA was extracted, purified, and quantified for reverse transcription using High Capacity RNA to cDNA Kit (Applied Biosystems) according to the manufacturer’s instructions. qPCR was carried out using 5 ng cDNA and 6 μl SYBR green master mix (Applied Biosystems). Each measurement was performed in triplicate.

#### Immunofluorescence and immunohistochemistry

iPSC-CMs were fixed in 4% paraformaldehyde for 5 min and blocked with 2% BSA for 1 hr. Cells were stained with Ki67 (1:5000, Abcam, ab11580) and TNNT2 (1:200, ThermoFisher, MS-295-P) at 4°C overnight. AlexaFluor-conjugated secondary antibody (Life Technologies) was then used and co-stained with DAPI (Vector Laboratories). Imaging was captured with a confocal microscope (Leica) and analyzed with ImageJ software.

The human control and HLHS heart paraffin sections were dewaxed, rehydrated and antigens were retrieved using Tris-based buffer (Vector Laboratories) for 20 min in the microwave. The samples were pre-treated with 0.5% H_2_O_2_ (Fisher Scientific) for 30 mins before blocked with 3% BSA (Sigma) for 1 hr. CDH11 (1:50, Invitrogen, 32-1700), CD31 (1:50, Abcam, ab9498), and FN1 (1:50, Abcam, ab6328) were used for overnight staining, followed by HRP-ABC development (Vector Laboratories) and hematoxylin (Vector Laboratories) nuclear staining. Imaging was captured using a bright field microscope (Leica) and analyzed using ImageJ software.

Frog embryos were fixed at stage 46 in the fixative MEMFA and dehydrated in ethanol and then in methanol overnight. The embryos were then rehydrated gradually into PBS and equilibrated in 20% sucrose solution. The embryos were next changed into OCT through graded washes, embedded in OCT, and sectioned at 12 microns. Sections through the heart was stained with anti-FN antibody (1:50, Developmental studies hybridoma bank-DSHB, 4H2) as previously described (Garmon et al., 2018). For FN signal intensity, a line was drawn across the ventricles and pixel intensities at the peaks of the plot were considered as FN signal at the inner lining of the ventricular chamber.

#### Angiogenesis assay

iPSC-ECs were starved in medium containing 0.2% FBS overnight and seeded at the density of 20,000 cells per 24-well plate in growth factor reduced Matrigel (Corning). The tube numbers were counted at 6 hours and 12 hours afterwards in three random fields. We defined a tube and quantified the number of tubes as described by Nickel et al. (Nickel et al., 2015).

#### BrdU proliferation assay

iPSC-ECs proliferation was assessed by using BrdU Cell proliferation kit according to the manufacturer’s instructions (Millipore). Minor modifications are as follows, 5000 cells in 100 μl EGM-2 were seeded per well in 96-well plate and cultured overnight. 20 μl of diluted BrdU label was added into each well, followed by a 24 hr incubation. The final results were subtracted from the background read.

#### Cell adhesion assay

Cell adhesion was carried out as previously described (de Jesus Perez et al., 2012). iPSC-ECs (10,000/well) were seeded on to a 24 well plate either uncoated (plastic) or coated with collagen IV (BD BioCoat^TM^), gelatin (0.2%, Sigma), laminin (BD BioCoat^TM^) or fibronectin (0.1 mg/ml, Sigma) and allowed to adhere for 1 hr. Non-adhesive cells were then washed away using PBS. The remaining adherent cells were stained with DAPI and imaged. The average number of cells was calculated by counting the total number of cells in six random views per well (10X magnification). All assays were repeated in triplicate.

#### Cell migration assay

The wound healing/cell migration assay was done using the Culture-insert 2 Well in u-Dish 35 mm system (Ibidi) following the manufacturer’s instructions. Briefly, 30,000 iPSC-ECs were seeded in each chamber and allowed grow overnight. After cell attachment, the Culture-insert 2 Well was gently removed using sterile tweezers. Then cells were washed with PBS and fresh EGM-2 culture medium was replaced. Migration was observed and imaged at 0, 9, and 24 hr.

#### Simulation of shear stress

μ-Slide I Luer (0.4 mm, Ibidi) slides were coated with 0.5% Gelatin for 30 min at 37°C. 75,000 iPSC-ECs in 100 μl EGM-2 were seeded onto the slides and cultured under static conditions for 24 hr. iPSC-ECs were then exposed to 15 dyne/cm^2^ of laminar shear stress (LSS) for 48 hours using the Ibidi Perfusion System (Ibidi). After the experiments, cells were either fixed with 4% PFA for imaging or subjected to RNA extraction using the RNeasy Plus Micro Kit (Qiagen).

#### siRNA transfection

siRNA transfection was carried out according to manufacturer’s instruction. ON-TARGET plus scramble, *ETS1*, *FN1*, *CHD7*, *KMT2D*, *FOXM1*, and *ARID1B* siRNA (GE Dharmacon) were transfected at 25 nM into subconfluent early passage iPSC-ECs or native endocardium using Lipofectamine RNAiMax (Invitrogen). 48 hrs after transfection, cells were either fixed for staining or subjected to RNA extraction.

#### Bulk RNA-Seq data analysis

The raw 150 bp paired-end reads were first trimmed as a quality control by TrimGalore v0.4.4 (https://www.bioinformatics.babraham.ac.uk/projects/trim_galore/) to exclude adapter sequences and bases with Phred scoring less than 20, reflecting a low probability of base-calling errors (p<1%), equivalent to >99% accuracy. Then, the trimmed pair-end reads were mapped to hg38 using STAR software(Dobin et al., 2013) with ENCODE options. FeatureCounts (Liao et al., 2014) v1.5.1 was used to count the number of reads in each gene annotated by Gencode (Harrow et al., 2012) v25. DESeq2 (Love et al., 2014) R package was used to detect DEGs between iPSC-CMs co-cultured with control or HLHS iPSC-ECs using the likelihood ratio test. Full model was set as full=∼treatment+1, and the reduced model was reduce=∼1. Genes with p-value<5% and fold change*≥*1.2 were define as DEGs. We used DESeq2 R package to normalize the raw counts from bulk RNA-seq data. The expression values of each candidate gene were extracted and followed by z-score normalization across all samples using “heatmap.2” function in R. The heatmap shows the z-score values of each gene in each sample. Gene Ontology (Ashburner et al., 2000) (GO) term enrichment analysis of differentially expressed genes was performed using the R package “GeneAnswers”(Lei Huang, 2018). GO terms with a false discovery rate (FDR)<5% were considered as significantly enriched. P value before the FDR correction for each functional annotated term was calculated from the hypergeometric test as follows:

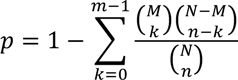

Where N is the total number of genes in any GO term; n is the number of differentially expressed genes in any GO term; M is the number of genes annotated for a particular GO term; m is the number of differentially expressed genes annotated for a particular GO term. Z score of a GO term was calculated as follows:

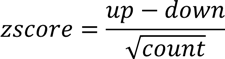

Where *count* is the total regulated gene number, *up* and *down* are the number of up-regulated or down-regulated differentially expressed genes, respectively. A custom R script was used for the data visualization.

### Single-cell RNA-Seq data analysis

#### Cell ranger for data pre-processing

Chromium Single Cell Software Suite v2.1.1 (https://support.10xgenomics.com/single-cell-gene-expression/software/pipelines/2.1/what-is-cell-ranger) was used for pre-processing the single cell RNA-seq data produced in the 10x Chromium Platform, which includes sample demultiplexing, read alignment, barcode processing, and UMI counting. “Cellranger mkfastq” was used to generate FASTQ files from BCL files. Specifically, the Illumina BCL output folder was used for sample demultiplexing, which is based on the 8 bp sample index read, and generating FASTQ files for the paired-end reads as well as the sample index by Illumina bcl2fastq. Only the first indexes were analyzed on the dual-indexed flowcell. Next, “cellranger count” was used to generate single-cell gene counts for a single library. Reads in the FASTQ files were mapped to the human reference genome (NCBI build38/UCSC hg38) with STAR software. Of note, for reads that aligned to a single exonic locus but also aligned to one or more non-exonic loci, only the exonic locus was prioritized. Read were considered to be confidently mapped to the exonic loci with MAPQ 255. Reads that mapped to the genome and transcriptome were delivered to a BAM file. Chromium cellular barcodes were used to generate gene-barcode matrices. Only reads that were confidently (uniquely) mapped to the transcriptome were used for the UMI count. Filtered gene-barcode matrices containing only cellular barcodes in the MEX format were used for downstream analysis.

#### Cellular clustering

The filtered gene-barcode matrix from the cell ranger pipeline for each sample was fed into the Seurat(Butler et al., 2018) R package for downstream analyses. Genes expressed in < 3 cells and cells with <500 detected genes or the percentage of mitochondrial genes >20% were removed. We used “MergeSeurat” command to combine samples. The raw read counts generated by cellranger were further normalized by a global-scaling normalization method, “LogNormalize”, that normalizes the gene expression measurements for each cell by the total read counts. The log-transformed normalized single cell expression values were used for visualizations (violin and feature plots) and differential expression tests. Variable genes with average expression in the interval [0.0125, 3], and dispersion*≥*0.5 were selected for Principal Component Analysis. Technical noise, batch effects and biological sources of variation such as number of UMI and the percentage of mitochondrial genes, were regressed to improve downstream dimensionality reduction and clustering. The scaled z-scored residuals were used for principal component analysis (PCA). The top 20 principal components were used for Uniform Manifold Approximation and Projection (UMAP). DEGs among clusters were detected by comparing cells in each cluster against all other cells using the Wilcoxon Rank Sum test. A gene is defined as a DEG in cluster X if it can be detected in *≥*25% cells in cluster X, with adjusted p-value<5% and fold change*≥*1.5 between cell in cluster X and all other cells. Methods for pathway enrichment analysis was the same as described in the bulk RNA-Seq data analysis. In the iPSC-EC pathway enrichment, only up-regulated genes in each cell cluster were selected, therefore no z score was expected. We followed the steps described in Paik et. Al(Paik et al., 2018) for cell-cell communication analysis. For each ligand-receptor pair, the mean expression of the ligand gene was calculated for all cells in the clusters classified as endocardium, and the mean expression of the receptor gene was calculated for all cells in the clusters classified as cardiomyocyte. The interaction value was estimated as the product of the two mean values: (mean expression of ligand gene in endocardial population) X (mean expression of receptor gene in myocardial population) and reflected as width of chords on circular plot using customized R package.

To generate the DNMs gene expression heatmap, we calculated their average expression values within one cluster, which were further normalized to z-scores across clusters using “heatmap.2” function in R. The heatmap shows the z-score values of each gene in each cluster.

### ChIP-Seq peak annotation and pathway analysis

ChIP-Seq peaks were downloaded from GSE93030, *ETS1* over expression induced *ETS1/RNAPII* binding peaks were generated by subtracting mod_GFP from mod_ETS1 in ETS1/ RNAPII ChIP-Seq peaks. Bedtools (https://bedtools.readthedocs.io/en/latest/) with -A parameter (i.e. remove entire peak if any overlap) was used for peak subtraction. Peaks were annotated using homer(Heinz et al., 2010) annotatePeaks.pl script set to the default parameters.

## QUANTIFICATION AND STATISTICAL ANALYSIS

First, normal distribution from each group was confirmed using *χ*^2^ test before any comparison between groups. Statistical analysis was then performed using Student’s *t*-test (two-sided) between two groups or ANOVA followed by Bonferroni post-test for multiple groups comparisons. If variances between two groups were significantly different (*F*-test), nonparametric Mann-Whitney test was applied. p<0.05 was considered as statistically significant. As for all the experiments, at least three independent experiments were performed unless otherwise specified. Analyses were carried out using GraphPad Prism 8.0 (GraphPad Software Inc., La Jolla, CA).

## Supplemental Figures

**Figure S1. scRNA-seq of CD144 positive cells from a normal human fetal heart (Related to Figure 1).** (A) UMAP projection of CD144 positive cells from day 83 normal human fetal heart in Figure 1B. UMAP projection of endocardial genes (B), valvular genes (C), arterial (D) and venous/capillary endothelial gene (E). Purple indicates high level of gene expression.

**Figure S2. Expression of endocardial and endothelial markers in iPSC-ECs and human fetal heart tissues from control and HLHS patients (Related to Figure 2).** (A) Gene expression levels of endocardial and endothelial markers on day 0 and day 8 of iPSC-EC differentiation determined by qPCR. n=3 in each group. Visualization of arterial and venous markers in control and HLHS iPSC-ECs by UMAP (B) and Violin plots (C). (D) qPCR detection of arterial and venous gene expression in iPSC-ECs from control (n=3) and HLHS (n=3) patients. (E) CD31 antibody authentication on normal fetal heart. CD31-labed pan-ECs were detected in the endocardium, coronary EC (endothelium), and capillary, as indicated by the black arrowheads. V: ventricular chamber; Co: coronary vessel; Ca: capillary. (F) CDH11 antibody authentication on normal fetal heart. Black arrowheads indicate positive staining. Of note, coronary EC (endothelium) was not labeled by CDH11, as showed by the red arrowhead. Data shown as the mean ± SEM. *p<0.05, control vs HLHS. In (A) and (D),

**Figure S3. Cell migration and flow response in control and HLHS iPSC-ECs (Related to Figure 3).** (A) iPSC-EC migratory rate at 9- and 24-hours post-scratch between control and HLHS patients. Yellow lines indicate the migration edges. Migration rate was normalized to 0 hour respectively. (B) Alignment of iPSC-ECs from control and HLHS patients under 48 hours, 15dyn/cm^2^ laminar shear stress (LSS). Arrow indicates the shear stress direction. (C) Shear stress related genes expression in iPSC-ECs under static or LSS conditions. Data shown as the mean ± SEM. *p<0.05, control (n=4) vs HLHS (n=3) under static condition.

**Figure S4. Bulk RNA-seq of normal control iPSC-CMs co-cultured with control or HLHS iPSC-ECs (Related to Figure 4).** (A) Heatmap of bulk RNA-seq DEGs in normal iPSC-CMs co-cultured with control or HLHS iPSC-ECs. Row z-score reflected the gene expression change. (B) KEGG pathway enrichment of cardiac muscle contraction related genes from bulk RNA-seq in iPSC-CMs co-culture with control or HLHS iPSC-ECs in Figure 4. Red defines up-regulation, green down-regulation.

**Figure S5. Single-cell RNA-seq comparing fetal heart with underdeveloped left ventricle vs. healthy control (Related to Figure 5).** (A) Schematic illustration for micro-dissection of day 84 human fetal heart with underdeveloped left ventricle (ULV). PA: pulmonary artery; RA: right atrium; LA: left atrium; RV: right ventricle; LV: left ventricle. (B) UMAP projection of various cell types from day 84 ULV. SMC: smooth muscle cell; RBC: red blood cell; CM: cardiomyocyte; EC: endothelial cell. (C) UMAP projection of represented genes for various cell types in human fetal heart. (D) UMAP projection of CD144 positive cells from day 84 ULV in (A). UMAP of endocardial genes (E), valvular genes (F), arterial and venous/capillary endothelial genes (G). (H) UMAP for scRNA-seq analysis comparing left ventricle ECs from control vs. ULV human fetal hearts. (I) UMAP projection of endocardial (I), endothelial (J), and pan-EC marker (K). (L) Functional enrichment analysis based on DEGs between ULV vs. Control endocardial populations. −log10FDR indicates the significance of enrichment.

**Figure S6. scRNA-seq analysis of iPSC-ECs and fetal heart ECs from control vs. patient, and isolation of cardiac EC subtypes from fetal hearts (Related to Figure 5).** (A) Analysis of overlapped DEGs between fetal heart endocardium (ULV vs. Control) and iPSC-EC endocardium (HLHS vs. Control). Monte-Carlo simulation calculated the overlapping significance of shared DEGs. (B) Gene name and fold change of shared DEGs from (A). (C) Immunostaining of FN1 protein in normal human heart showed positive labeling of coronary endothelium (Left), ventricular endocardium (Middle), and atrial endocardium (Right) as indicated by black arrowheads. Asterisks indicate small coronary vessel/capillary with low expression of FN1. (D) Work flow for isolation of human fetal endocardium, coronary endothelium, and aortic endothelium. (E) qPCR of markers for different cardiac cell types in isolated human fetal heart. Primary cardiac fibroblast and iPSC-CMs were used as positive controls. EC: CD144^+^-sorted cells; non-EC: CD144 negative cells. Data shown as the mean ± SEM. **p<0.01, EC vs non-EC. (F) qPCR verification of the purity of different cardiac EC subtypes isolated from human fetal heart. Data shown as the mean ± SEM. **p<0.01, coronary endothelium vs. endocardium; ##p<0.01, aortic endothelium vs. endocardium. n=3 in each group.

**Figure S7. Transcription factors and chromatin remodelers associated with HLHS DNMs control endocardial gene expressions (Related to Figure 6).** (A) Expression levels of the genes harboring HLHS DNMs were reduced in HLHS iPSC-ECs compared with controls. Control (n=4) vs HLHS (n=3). Expression of *FN1*, *NPR3*, and *CDH11* in native endocardial cells with knock-down of *KMT2D* (B), *ARID1B* (C), or *FOXM1* (D), respectively. (E) Top: RNAPII and histone marker ChIP-seq tracks around *CDH11* gene in HUVECs. The black box and red shading indicate the designated location of primers for ChIP-qPCR. The red arrow indicates the transcription start site (TSS) direction. Bottom: ChIP-qPCR of ETS1 or CHD7 binding to *CDH11* enhancer regions in control and HLHS iPSC-ECs. (F) *ETS1* UMAP projection and violin plot of iPSC-ECs from control and HLHS patients. (G) ChIP-qPCR detecting RNA Pol II binding capacity to the promoter regions of endocardial genes *NPR3*, *PLVAP*, *CDH11* and *FN1* in the endocardium after *ETS1* knock-down (H) ChIP-seq track of *NPR3* and *PLVAP* promoter regions, indicated by H3K27ac and H3K4me3 peaks, from HUVECs retrieved either from ENCODE (RNAPII, H3K4me3, and H3K27ac) or Chen et al. (Mod_GFP and Mod_ETS1). Data shown as the mean ± SEM. *p<0.05, **p<0.01, control (n=3) vs HLHS (n=3) or scramble (n=3) vs siRNA knock-down (n=3).

