## Supplementary figures and images for "Single-Cell RNA-Seq Reveals Endocardial Defect in Hypoplastic Left Heart Syndrome"

### Supplemental Figures

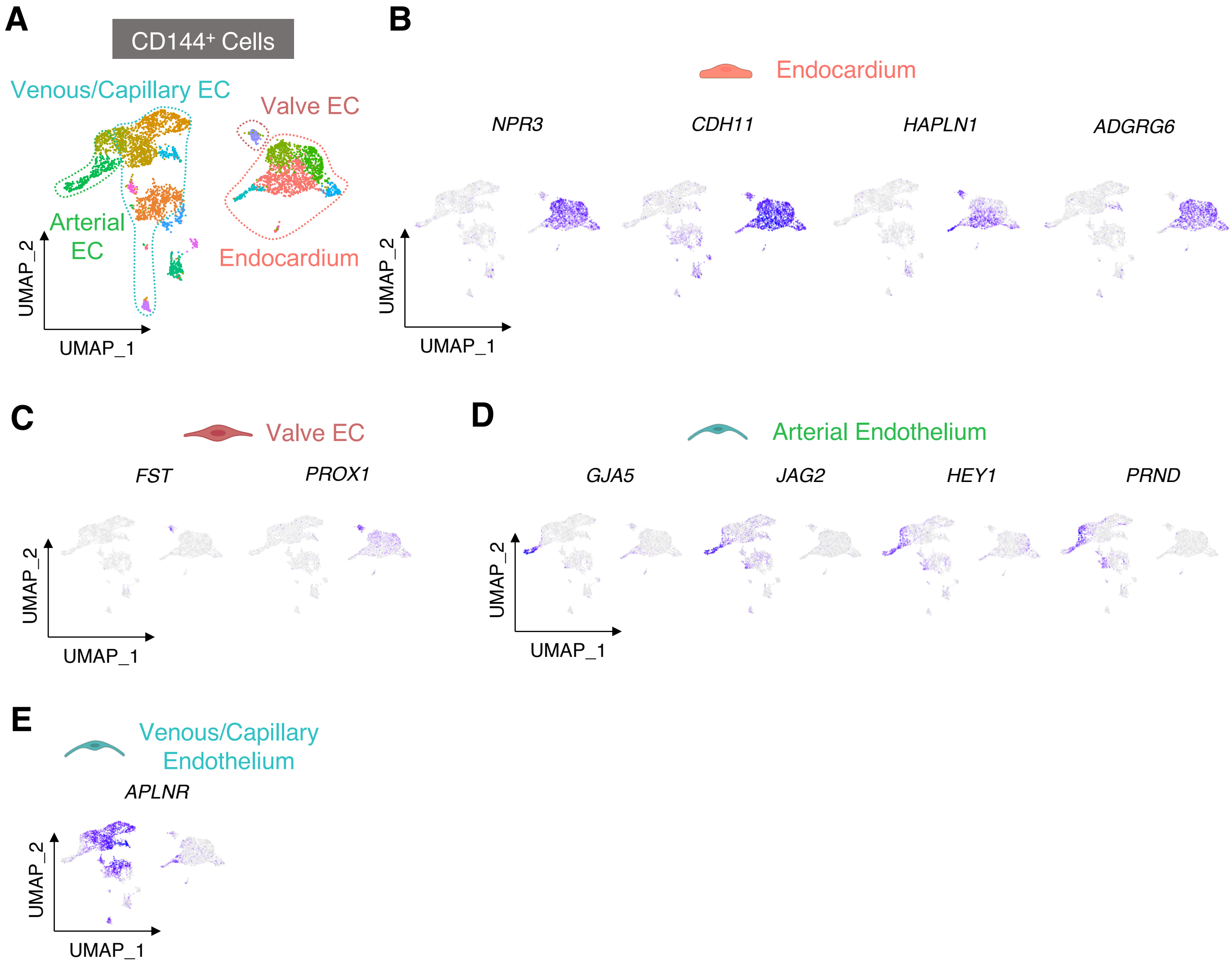

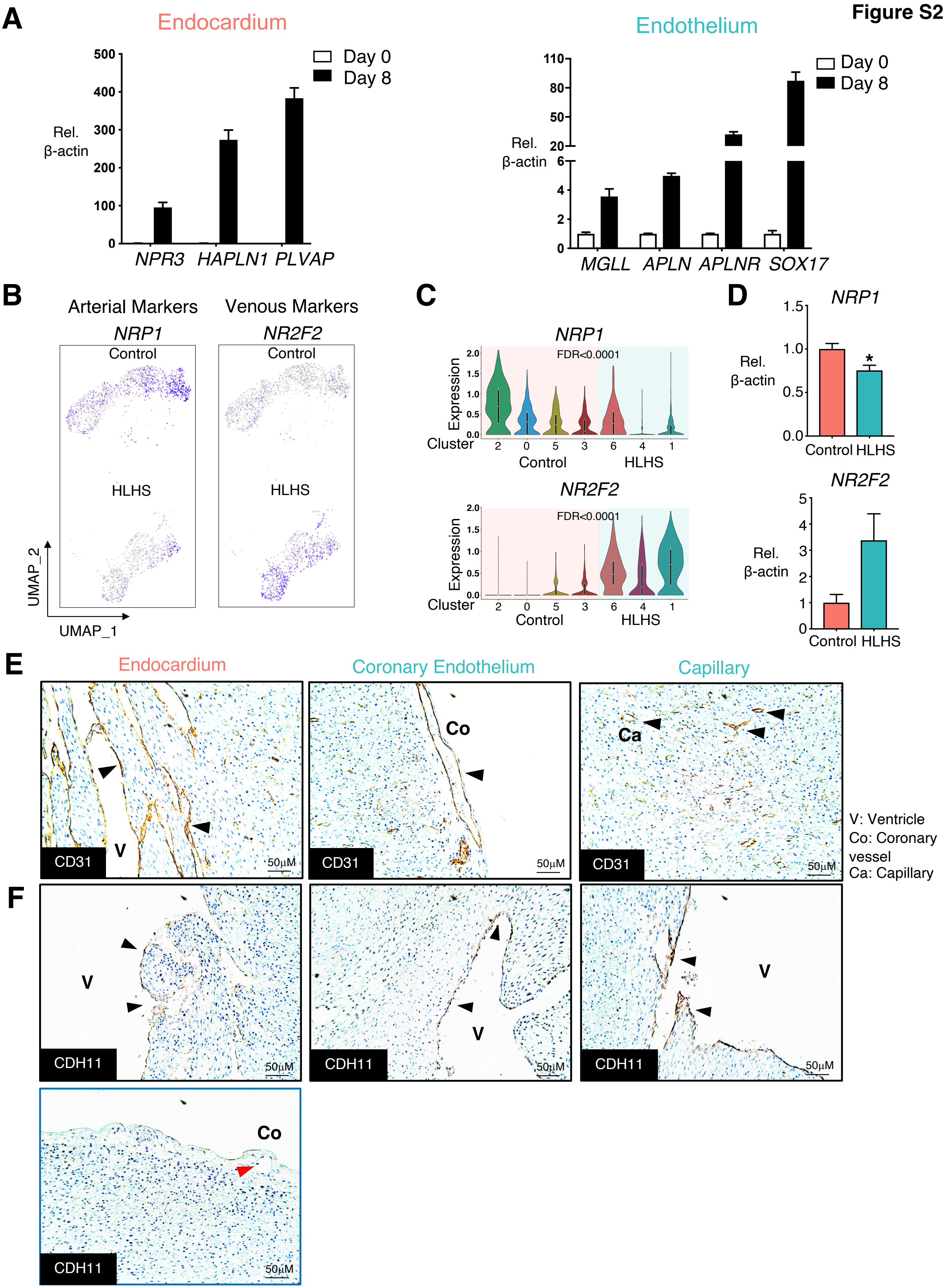

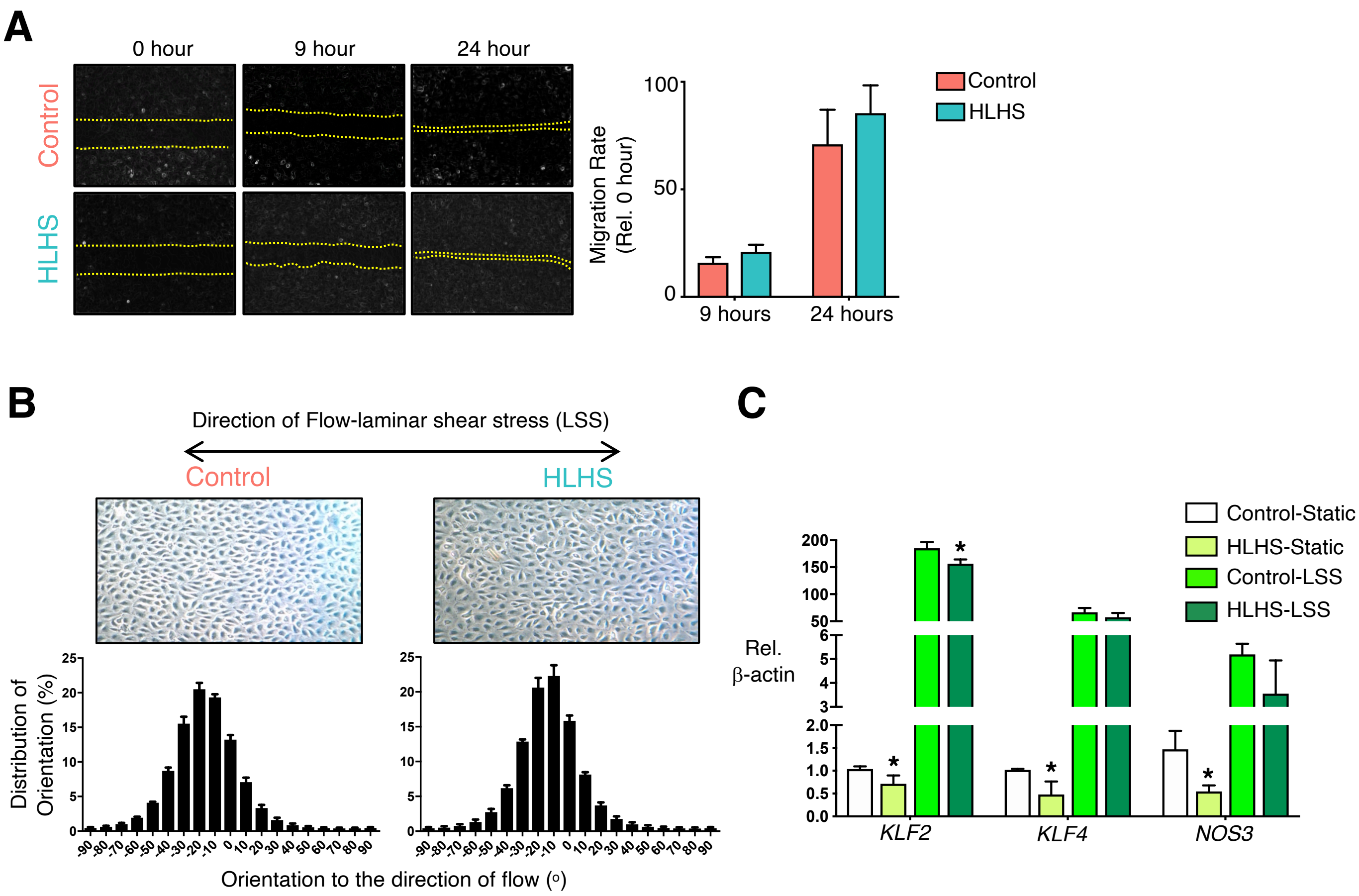

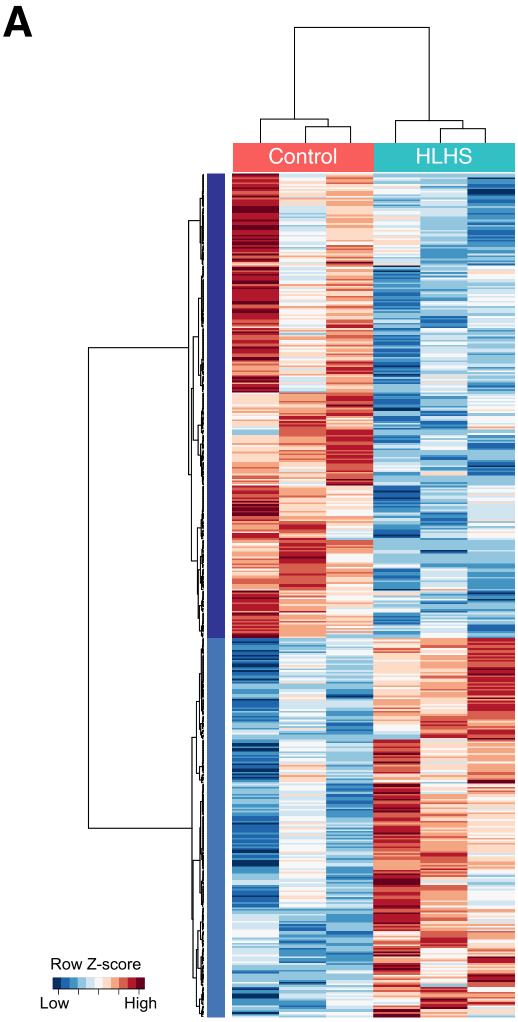

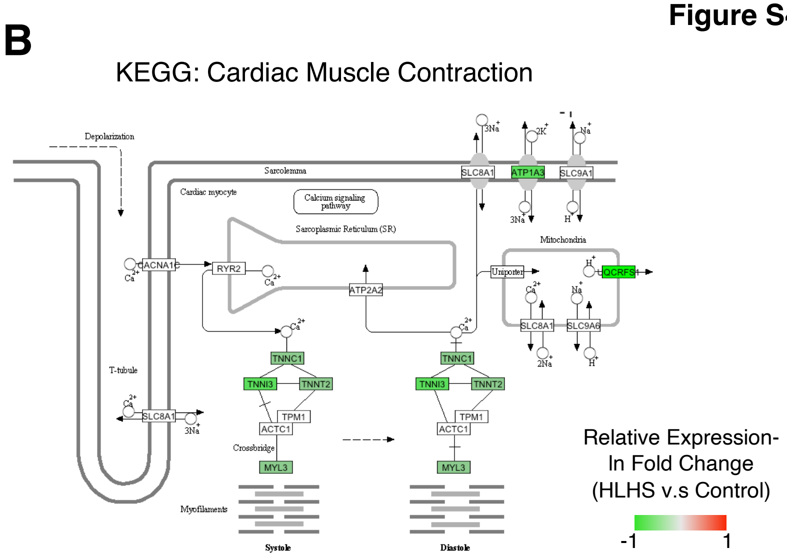

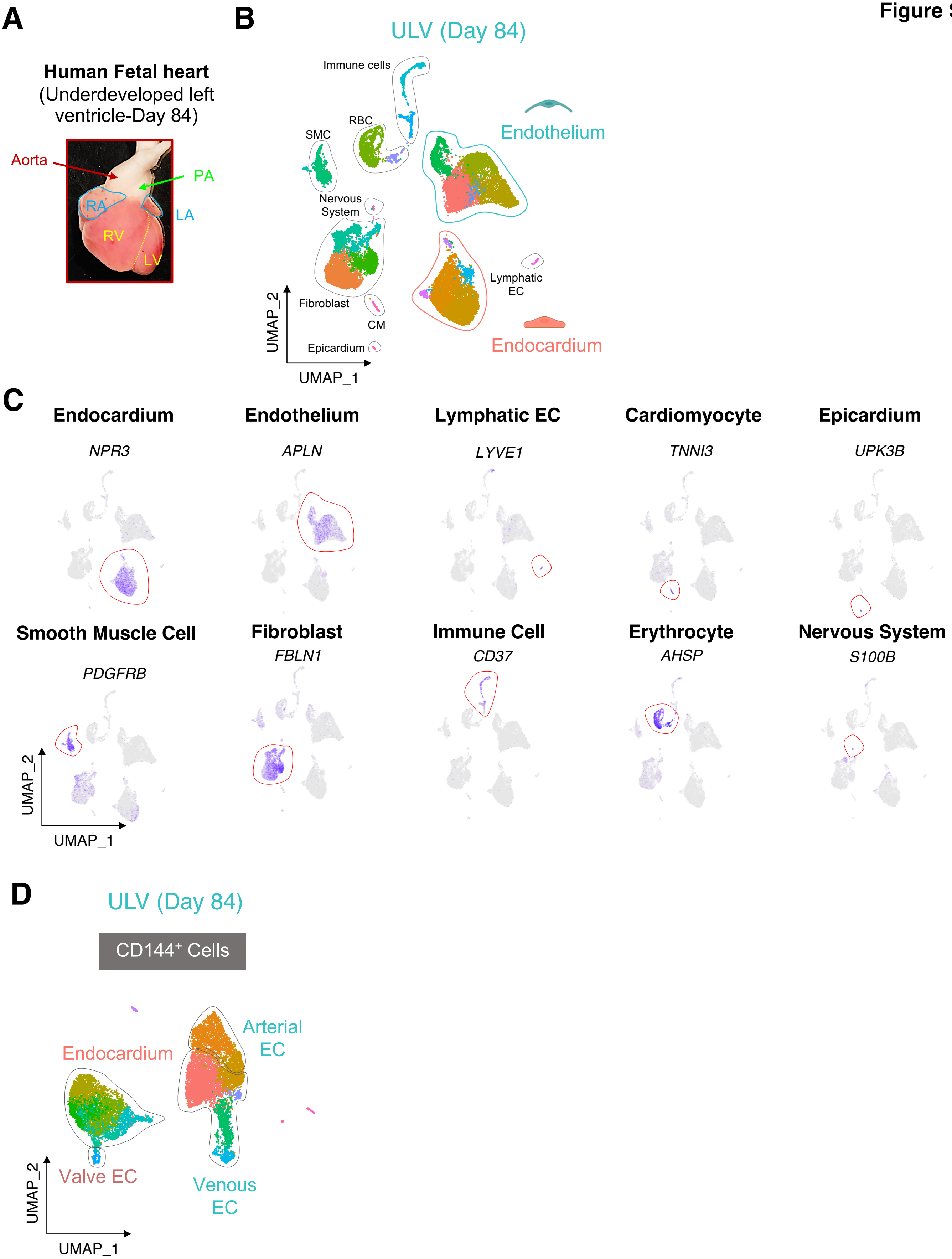

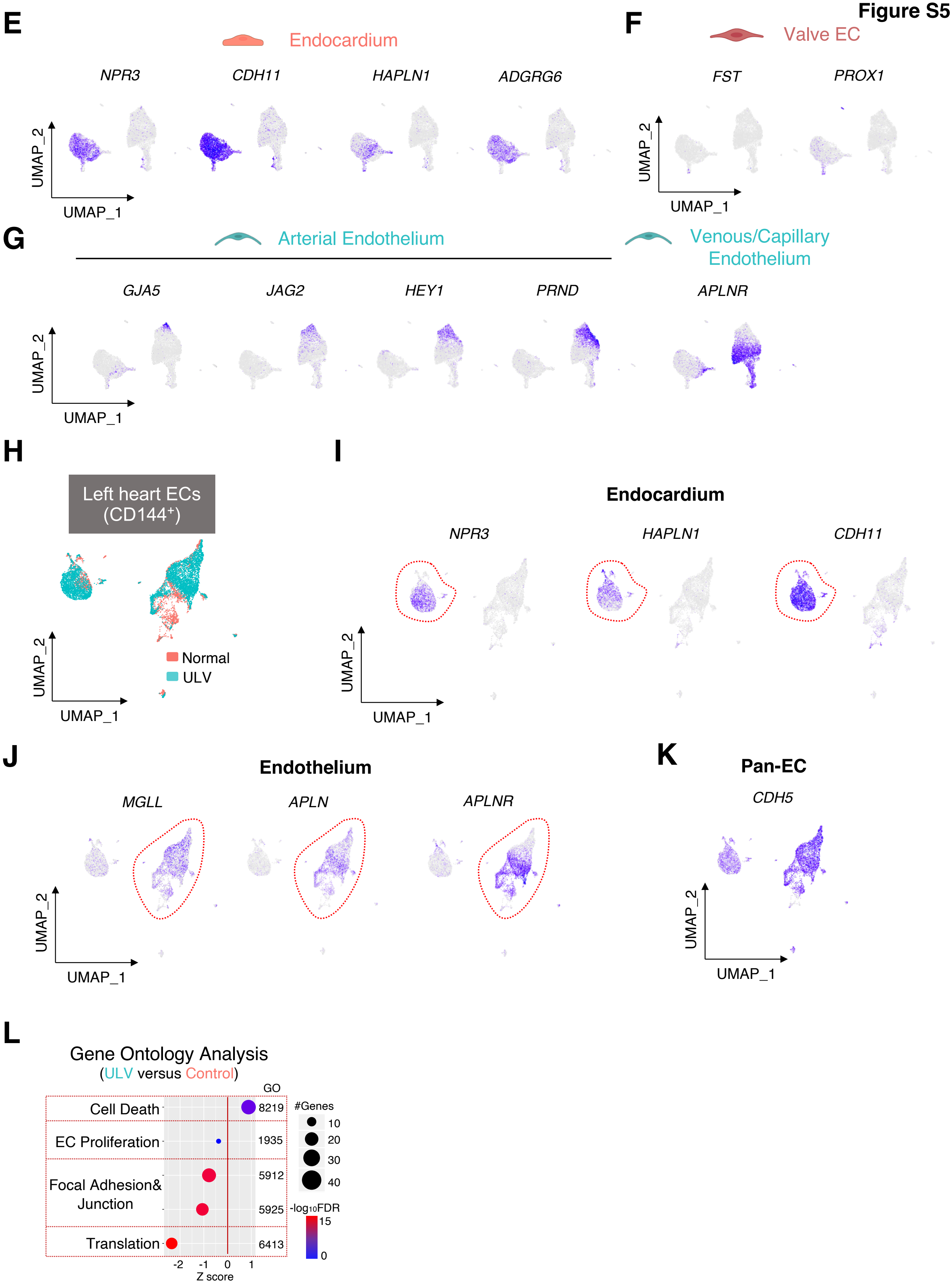

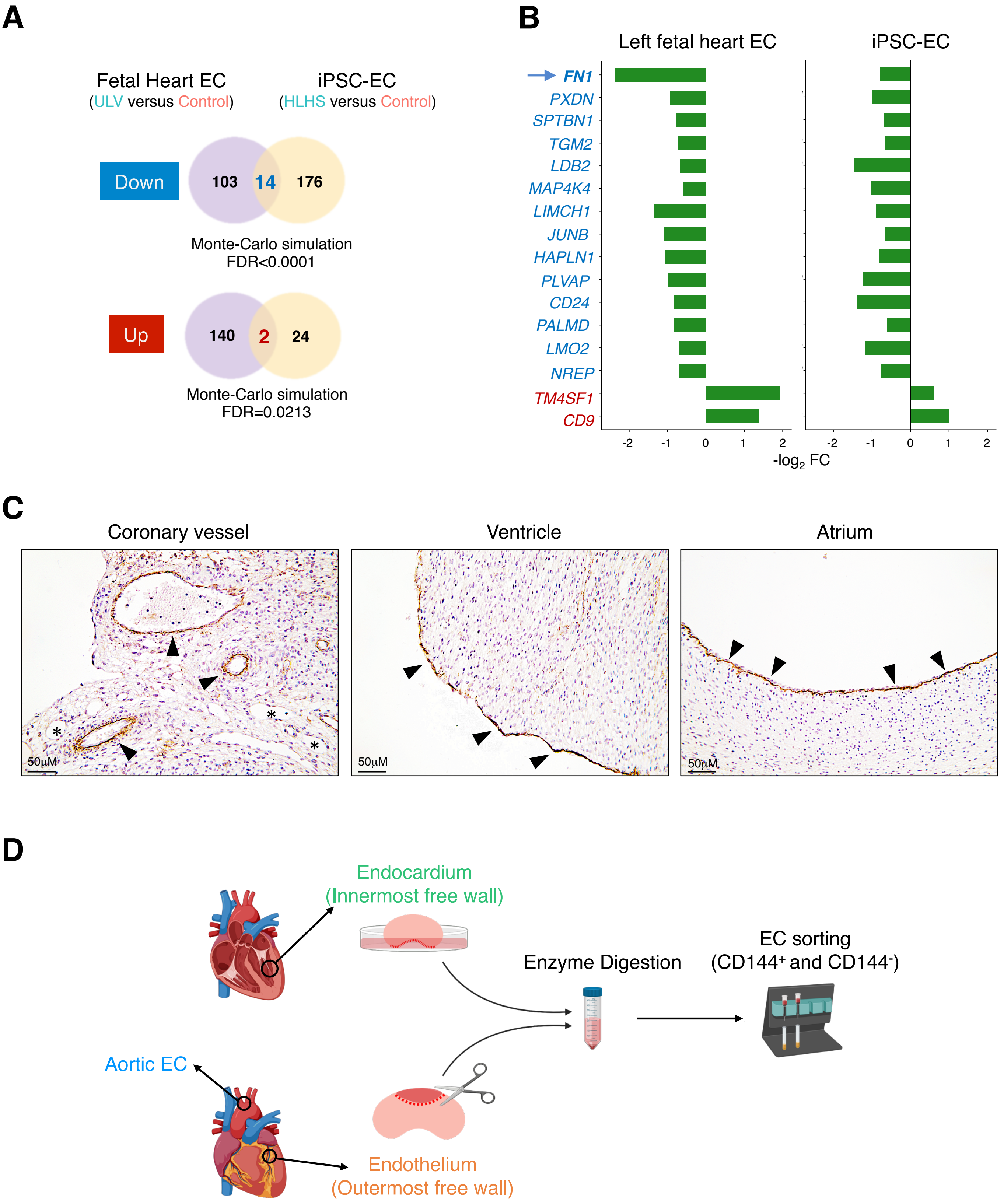

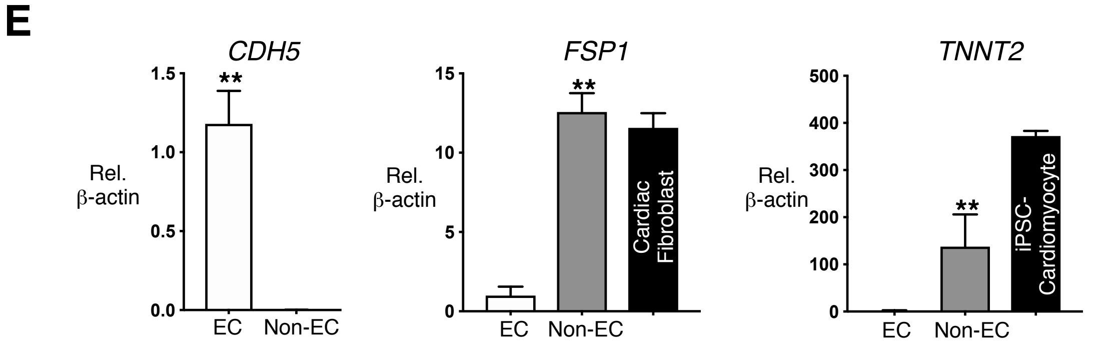

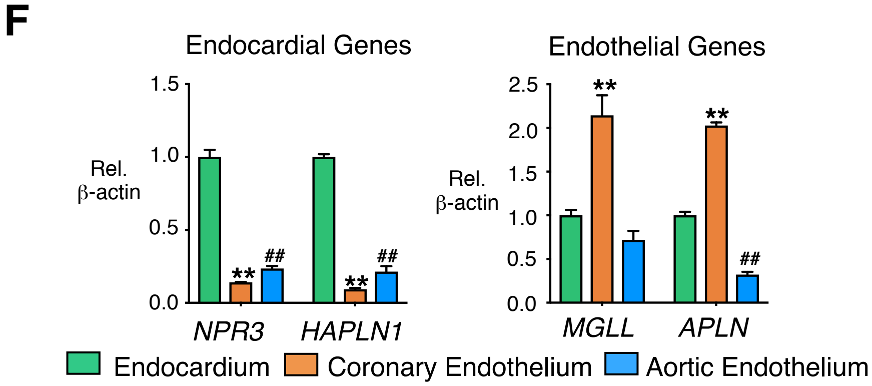

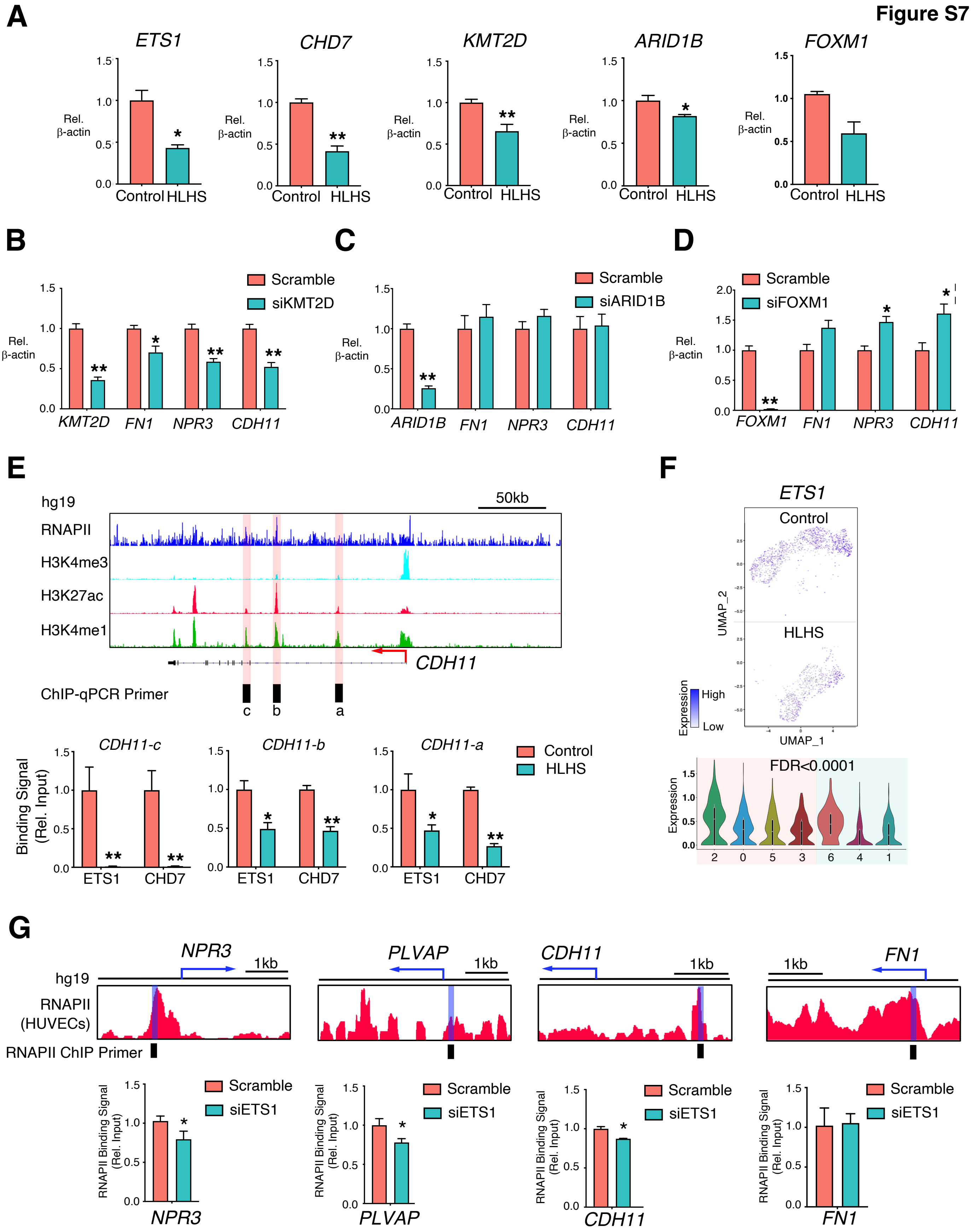

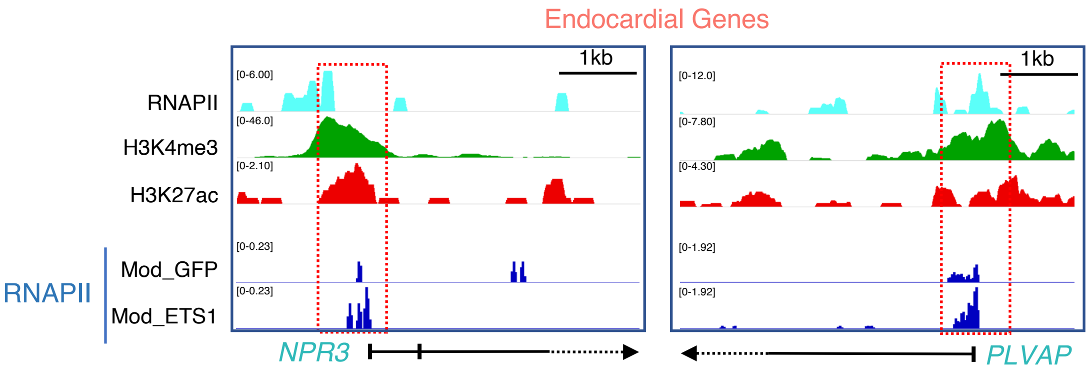
